# Impact of Prescribed Fire on Soil Microbial Communities in a Southern Appalachian Forest Clearcut

**DOI:** 10.1101/2024.04.09.588751

**Authors:** S.A.A Rafie, L. Blentlinger, A. D. Putt, D. E. Williams, D. C. Joyner, M. F. Campa, M. J. Schubert, K. P. Hoyt, S. P. Horn, J. A. Franklin, T. C. Hazen

**Affiliations:** Department of Civil and Environmental Engineering, University of Tennessee, Knoxville, TN, USA 37916; Department of Geography, University of Tennessee, Knoxville TN, USA 37916; Department of Forestry, Wildlife and Fisheries, University of Tennessee, Knoxville, TN, USA 37916; Bredesen Center – Genome Science & Technology, University of Tennessee, Knoxville, USA 37916; Institute for a Secure and Sustainable Environment, University of Tennessee, Knoxville, TN, USA 37916; □Oak Ridge National Laboratory Oak Ridge, TN, USA 37831; Department of Earth & Planetary Sciences, University of Tennessee, Knoxville, TN, USA 37916; Forest Resources AgResearch and Education Center, Knoxville, TN, USA 37916

**Keywords:** prescribed fire, microbial diversity, soil recovery, soil microbiome, nutrient cycling

## Abstract

Escalating wildfire frequency and severity, exacerbated by shifting climate patterns, pose significant ecological and economic challenges. Prescribed burns, a common forest management tool, aim to mitigate wildfire risks and protect biodiversity. Nevertheless, understanding the impact of prescribed burns on soil and microbial communities in temperate mixed forests, considering temporal dynamics and slash fuel types, remains crucial. Our study, conducted at the University of Tennessee Forest Resources AgResearch and Education Center in Oak Ridge, TN, employed controlled burns across various treatments, and the findings indicate that low-intensity prescribed burns have none or minimal short-term effects on soil parameters but may alter soil nutrient concentrations, as evidenced by significant changes in porewater acetate, formate, and nitrate concentrations. These burns also induce shifts in microbial community structure and diversity, with Proteobacteria and Acidobacteria increasing significantly post-fire, possibly aiding soil recovery. In contrast, Verrucomicrobia showed a notable decrease over time, and other specific microbial taxa correlated with soil pH, porewater nitrate, ammonium, and phosphate concentrations. Our research contributes to understanding the intricate relationships between prescribed fire, soil dynamics, and microbial responses in temperate mixed forests in the Southern Appalachian Region, which is valuable for informed land management practices in the face of evolving environmental challenges.

## 1. Introduction

The frequency and severity of wildfires are expected to increase with the changing climate, which can have significant ecological and economic consequences. Prescribed burns are commonly used in grassland and forest management to reduce the risk of wildfires and maintain wildlife habitats, particularly in the Southern Appalachian region (Lafon et al., 2017). However, the occurrence of large, severe wildfires during droughts in the region highlights the need to understand the effects of burning on the biogeochemical characteristics of soils. Over the past three decades, the extent of wildfire destruction has surged by a factor of four, with a concomitant rise in high-severity fires documented by the Monitoring Trends in Burn Severity (MTBS Project) study (MTBS Project, 2022). This trend is notably prominent in the Southeastern region of the United States, which has experienced a substantial expansion in prescribed burn areas compared to relatively stable conditions in other regions, as outlined in a study by Burke and collaborators (Burke et al., 2021).

Numerous studies have examined the causes of fire events in the Southern Appalachian region. One key factor is that the region is characterized by hot, humid summers and mild, wet winters. This climate creates favorable wildfire conditions, particularly during drought (Wear and Greis, 2013). Human activities, such as campfires and intentional burning, can also contribute to regional fire events (Abatzoglou and Williams, 2016). In terms of the ecological effects, wildfires can have both positive and negative impacts. It is widely accepted that wildfires can be essential for maintaining the diversity of plant and animal species in forest ecosystems and, as such, play a key role in the maintenance of ecosystem services (Coogan et al., 2019). Ecosystems like grasslands and savannas are fire-dependent and rely on recurring burns to preserve the characteristic structures, highlighting the positive impact of fires on soil health (Pivello et al., 2021). However, the increased frequency of fire events has undesirable consequences on human health, for example, by exacerbating pollutant and treatment by-product concentrations in nearby water sources (Paul et al., 2022). Wildfires can increase erosion and sedimentation rates, impacting water quality and aquatic habitats (Sankey et al., 2017). Air quality in the US has been affected by the spike in fine particulate matter (PM_2.5_) for extended periods following large-scale fire events over the past decade and exposed millions of people to harmful air quality (Jaffe et al., 2020). The economic and social impacts of wildfires in the Southern Appalachian region have also been studied extensively. These impacts can be significant, particularly for communities that rely on tourism or agriculture, since wildfires can damage or destroy homes, businesses, and other infrastructure, inflicting long-lasting economic consequences (Paveglio et al., 2016). Studies have shown that management practices can help reduce the severity of wildfires in the Southern Appalachian region (Hiers et al., 2016).

Prescribed burning can reduce the fuel available for fires, lowering the risk of severe wildfires. Other management practices, such as thinning and fuel reduction, have also been effective (Waldrop et al., 2010).

The extent and severity of fire can vary depending on the fuel type and burn history, and these factors can impact the rate and trajectory of post-fire soil recovery. Understanding these relationships is critical for predicting the long-term effects of wildfires on ecosystem function and resilience. Fires in forests dominated by coniferous trees can lead to more significant soil erosion and nutrient loss than in mixed or deciduous forests (Neary et al., 1999; Certini, 2005; Moody and Kinner, 2006). Coniferous forests tend to have less protective ground cover or exposed mineral soil, which can lead to greater soil disturbance during a fire. The burn history of an area can also affect post-fire soil recovery. Repeated burning can change soil organic matter content, nutrient availability, and microbial communities (Fonturbel et al., 2021). These changes can impact the trajectory and rate of post-fire soil recovery. Areas that have experienced frequent fires may have lower soil organic matter content, leading to lower soil fertility and slower recovery rates.

The interaction between fuel type and burn history in post-fire soil recovery is complex, especially when considering regions with varying fire histories (Keeley and Syphard, 2019). Recent changes in climate appear to have increased the frequency and severity of droughts and fires in various regions of North America. The impacts of fires on nutrient cycling in surface soils can vary depending on the percentage of fuels consumed. For instance, post-fire nitrogen (N) content in the soil may considerably decrease due to the loss of fuel N amounts, while short-term increases in soil ammonium (NH_4_ ^+^) and nitrate (NO_3_ ^−^) have also been observed (Knoepp and Swank, 1993; Certini, 2005; Ramirez et al., 2010; Taylor and Midgley, 2018). While low-severity fires can facilitate the rapid uptake of nutrients by intact roots to produce new above-ground biomass, greater fire severity may cause surface soils to reach lethal temperatures, temporarily reducing the influence of soil biota on nutrient fluxes. Moreover, the changes in soil properties and nutrient bioavailability brought about by fires can also have short- and long-term effects on the structure and activity of the soil microbial community (Schimel and Schaeffer, 2012; Alcaniz et al., 2018; Huffman and Madritch, 2018; Fischer et al., 2023). A reduction in microbial biomass due to heat is frequently observed in the soil surface layer (0–5 cm) following a fire (Alcaniz et al., 2018; Barreiro and Diaz-Ravina, 2021).

The influence of fire on soil properties and microbial communities, intricately linked to burn severity, has been the focus of previous research, as highlighted by (Adkins and Miesel, 2021). However, within the context of temperate mixed forests, there is a notable deficiency in our understanding of how prescribed fire events impact the functional potential and metabolic capacity of the soil microbiome. Studying microbial responses to environmental stressors in extreme environments offers valuable insights into microbial adaptability and emphasizes the role of microbial diversity and metabolic capabilities in ecosystem resilience (Chivian et al., 2008; Hemme et al., 2010).

Research on microbial responses concentrates on soils within specific forest biomes, such as Boreal or Chapparal forest soils (Mataix-Solera et al., 2009; Tas et al., 2014; Adkins and Miesel, 2021). While the effects of fire on soil properties and microbial communities have been extensively studied in these ecosystems, there remains a substantial gap in our understanding of these intricate relationships within the context of temperate mixed forests. These unique ecosystems, with their distinctive characteristics, call for a more comprehensive exploration of the impacts of prescribed fire, soil dynamics, and microbial responses. This study aims to investigate the impacts of prescribed fire on soil properties, microbial communities, and their interactions in a temperate mixed forest in the Southern Appalachian region. It focuses explicitly on how time and slash fuel types influence these effects. Our hypotheses are as follows: (i) prescribed burns will result in minimal impacts on overall soil chemistry but may lead to changes in soil nutrient concentrations; (Bolyen et al.) prescribed burns will induce alterations in microbial community structure and diversity, potentially enhancing functional redundancy within soil bacterial communities; and (iii) bacterial community composition is correlated to changes in soil nutrients.

## 2. Materials and methods

### 2.1. Site Description

The study site (Figure 1) is located in the temperate deciduous forest region at the University of Tennessee Forest Resources AgResearch and Education Center (FRREC) in Oak Ridge, TN, USA (GPS: 36° 1’ 12.08” N, 84° 11’ 35.47” W). It was established in 2017 on a former clear-cut site. Before cutting, the site was a mixed hardwood forest (52% white oak, 36% red oak, 8% poplar, 5% other hardwoods, and <1% pine). Controlled burns were carried out in late Fall (November) 2017 and Spring (March) 2019 across three treatments (no added fuel, added hardwood, and added pine) in triplicate blocks (for example, 1-1, 1-2, 1-3; for a more detailed site map please see Figure S1 of supplementary materials). “No-burn” plots were also established and serve as control. The quarter-acre (0.1 hectare) plots were subjected to controlled burns using the ring-fire technique(Wade and Lundsford, 1990), while fire intensity was monitored with a thermal imaging (FLIR) camera. These controlled burns resulted in very low-intensity fires, with patches of charred slash remains.

**Figure 1.**
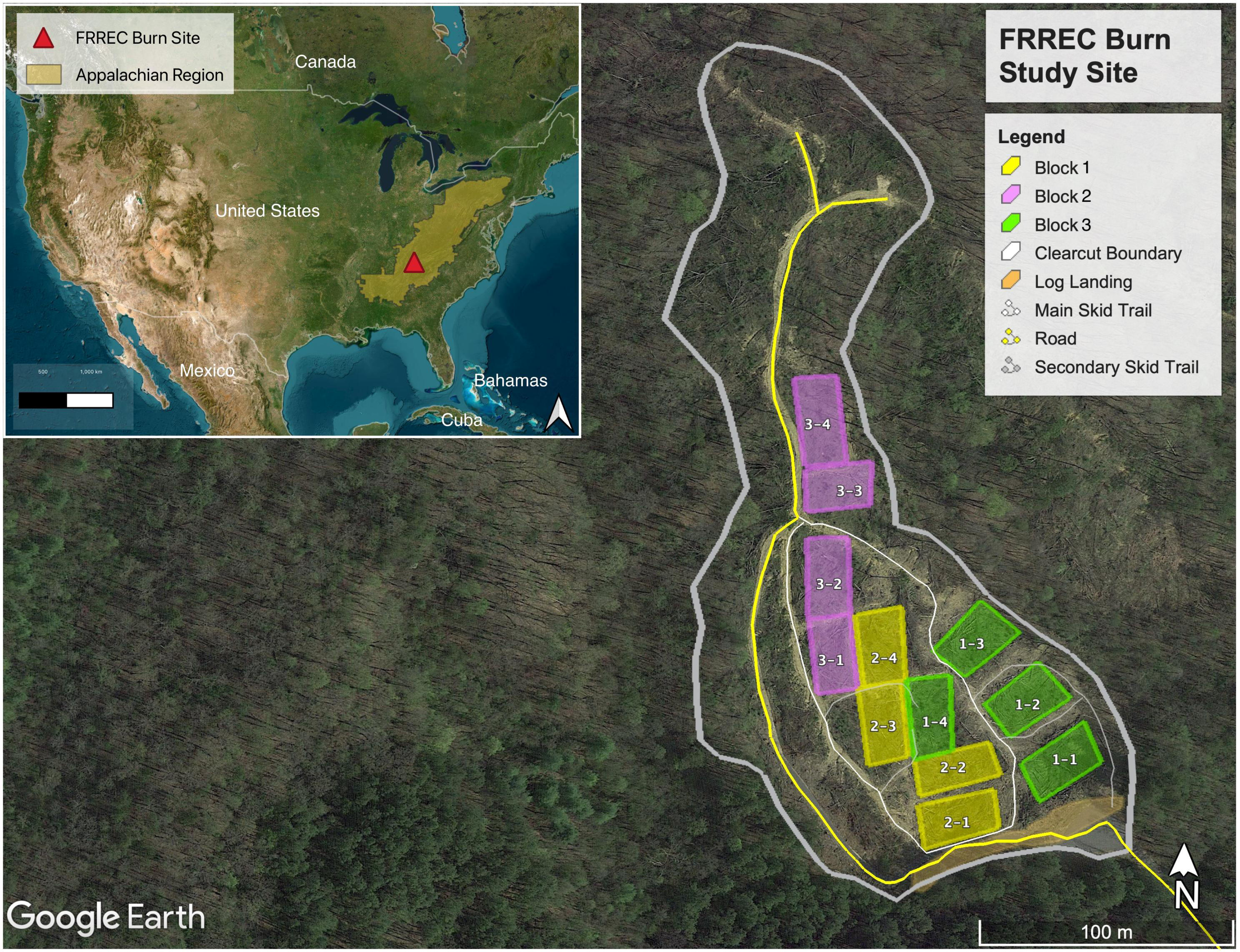
Map of FREEC burn sites (with United States and Appalachian region shown in inset) with fuel loading. Three treatments (no added fuel, added hardwood, and added pine) were set up in triplicate blocks (color-coded). “No-fire” control plots can be the fourth treatment (the last plot of every block). See Figure S1 in Supplementary Materials for more details.

### 2.2. Sampling

Samples were collected at four different time points: T0 - Pre-Burn, T1 - Immediately After Burn, T2 - 3 Months After Burn, and T3 - 17 Months after Burn. Five samples were collected from each plot for T0 and T1, and triplicate sampling for T2 and T3, with the locations being randomly chosen (from generated random points based on what can be measured from the central post). We collected the top 5 cm of soil, which included both O and A horizons. Samples were cored out using sterile and truncated 50 ml syringes, which were immediately put on dry ice and then stored at −80°C until extraction was needed. These samples were subsequently used for microbial and soil geochemical analyses.

### 2.3. Soil Geochemistry

pH analysis were conducted following (Kalra and Maynard, 1991) with adjustments made according to (Miller and Kissel, 2010). Soil pH in 0.01 M CaCl_2_ was determined using a 1:1 soil-to-solution ratio and an electronic pH meter. Anions and organic acids were measured on pore water sample collected by filtration through 0.22 μm pore-sized filters. The filtered samples were then diluted and stored in 1.0 ml vials for subsequent analysis. Concentrations of anions were determined using a Dionex™ ICS 5000+ series (Thermo Fisher Scientific, Waltham, MA, USA) equipped with an AS11HC column operated at 35°C, with a KOH effluent gradient ranging from 0 to 60 mM at a flow rate of 1.3 ml/min. The working range for anion concentrations was set at 0.1-200 mg/l. Likewise, organic acids were quantified using the same system with a working range of 5-200 μM.

Cations were determined using a Dionex™ ICS 5000+ series with a CS12-A column at 35°C, using an isocratic 20 mM methanesulfonic acid effluent at a flow rate of 1 ml/min. The working range for cation concentrations was 5 μg/l - 500 mg/l. Before analysis, samples were acidified with a 10-vol% 1 M HCl solution to ensure the stability of the analytes.

### 2.4. Genomic DNA extraction and PCR amplification

DNA Extraction was carried out using the DNeasy® PowerSoil® Pro Kit (Qiagen) with 250mg of homogenized soil from each sample. This study used a DNA amplification approach, as described by supplementary methods in (Caporaso et al., 2012). The V4 region of the 16S rRNA gene was amplified together with barcodes (Illumina, San Diego, CA, USA) adaptor and sequencing primers, and the target primers, 515F and 806R. The efficiency of the amplification process was assessed through agarose gel electrophoresis. The PCR products were then pooled for each amplicon based on gel electrophoresis band strength and combined in equimolar concentrations.

### 2.5. 16S rRNA gene amplicon sequencing

Purification of PCR products was carried out using a DNA Clean & Concentrator Kit (Zymo). PCR products were checked for quality and quantity with the Agilent Bioanalyzer 2100. Based on Bioanalyzer results, Sequencing library preparation was conducted per the guidelines provided in the MiSeq™ Reagent Kit Preparation Guide (Illumina, San Diego, CA, USA) as outlined by (Caporaso et al., 2012). The sequencing process involved forward, index, and reverse reads over 251, 12, and 251 cycles. Sequencing was performed on the Illumina MiSeq platform, utilizing two separate 300-cycle v2 Illumina MiSeq kits to accommodate all samples.

Sequencing analysis was performed using Qiime2 (version 2020.11) (Bolyen et al., 2019). FASTQ files from the two separate sequencing runs were imported and then demultiplexed, followed by removing forward and reverse primers using the q2_cutadapt plugin in Qiime2. The DADA2 plugin (Callahan et al., 2016) with a paired-end setting was utilized to denoise and eliminate chimeric sequences and low-quality regions to generate Amplicon Sequence Variants (ASVs). The first 13 bases of each forward and reverse sequence were trimmed off for quality filtering, and each paired read was truncated at 150 bases. Representative 16S ASVs were assigned using SILVA 132 pre-trained gene databases (Quast et al., 2013). Sequences assigned to mitochondria and chloroplast for bacteria were removed from the ASV tables before subsequent analysis. Microeco (Liu et al., 2021), Vegan (Oksanen et al., 2016), and phyloseq (McMurdie and Holmes, 2013) packages were used in R (R Core Team, 2021) for community analysis. The trans_func class in the microeco package was used to match the taxonomic information of prokaryotes against the FAPROTAX(Louca et al., 2016) database for the functional annotation of prokaryotic taxa. This entailed establishing correlations between ASV abundances and traits from the database alongside environmental factors. Subsequently, this data was utilized to conduct differential tests on ASV abundances at the phylum level associated with specific traits across various groups. Raw reads used in this study can be found in the NCBI Sequence Read Archive (SRA) under the BioProject accession number PRJNA1027882.

### 2.6. Statistical analyses

All statistical analyses were performed using R v.4.2.3 software (R Core Team, 2021). The two Illumina runs resulted in 3.1 M and 13.8 M bacterial sequences, averaging 201,400 sequences/sample. We took a conservative approach and accounted for uneven sequencing depth by rarefying ASV tables to 5,022 sequences/sample depth (rarefaction curves can be found in Supplementary Materials, Figures S2 and S3). Using this approach, we retained the largest number of samples and sequencing depth within each dataset. Kruskal-Wallis (KW) tests followed by Dunn’s test for multiple comparisons across groups were performed to determine any significant changes in soil properties over time. Principal Coordinate Analysis (PCoA) was performed to explore the relationships between soil variables (pH, total C, total N, C: N ratio, porewater cations, and organic acids). We ran differential abundance tests using differential expression tools for compositional data (using ALDEx2 or ANOVA) through the trans_diff class in the microeco package. The comparisons were not restricted to pairs and were used to show whether there were any statistically significant changes (p <0.05) in the relative abundance (centered log-ratio normalized) for the top ten most abundant ASVs across time and different fuel types. The comparisons were followed by a Duncan’s multiple range post hoc test. Beta diversity analyses included non-metric multidimensional scaling (NMDS) and PcoA visualizations based on the weighted-Unifrac distance metrics. Nonparametric permutational multivariate analysis of variance (PERMANOVA) was performed using the weighted Unifrac distance matrix at a phylum level within the “vegan” package (Oksanen et al., 2016). Redundancy analysis was performed to determine which environmental variables influenced changes in microbial community composition using the trans_env function of the microeco package and passed to the “rda” function of vegan package to normalize the taxa data. Environmental variables were center log-ratio (CLR) normalized using the decostand function in “vegan” prior to input. RDA ordination was performed using the weighted Unifrac distances, and we used ANOVA through the cal_ordination_anova() and cal_ordination_envfit() functions to check the significance of the ordination model. The cal_mantel() function in microeco was used to apply a Mantel test and check for significant correlations between environmental variables and distance matrix. Correlation heatmap was generated using relative abundance data at Genus level with the cal_cor function with p-value adjustments (Benjamini & Hochberg method) performed for each environmental variable separately.

## 3. Results

### 3.1. Physicochemical properties of soil

Figure 2 shows the variations observed at different time points for porewater concentration of select cations, anions, and soil parameters like pH and Total Carbon and Nitrogen content. Porewater chemistry showed minimal changes in post-burn samples, with minor differences across fuel types or collection time. Significant increases (p < 0.05, KW-Dunn; See Tables T1 and T2 in supplementary materials) in acetate, formate, and nitrate concentrations and pH indicated changes in solute chemistry post-fire. The bulk carbon concentration in the soil cores exhibited a wide range, spanning from 1.23% to 6.98%. Analyzing the soil cores’ total carbon content (% wt.) revealed that in Pine Slash and No Added Slash treatments, showed increased total carbon content following the burn event. In contrast, when measured 17 months later, there was a notable decrease in total carbon content for these treatments. Due to the low-intensity burn, very few soil parameters were significantly affected.

**Figure 2.**
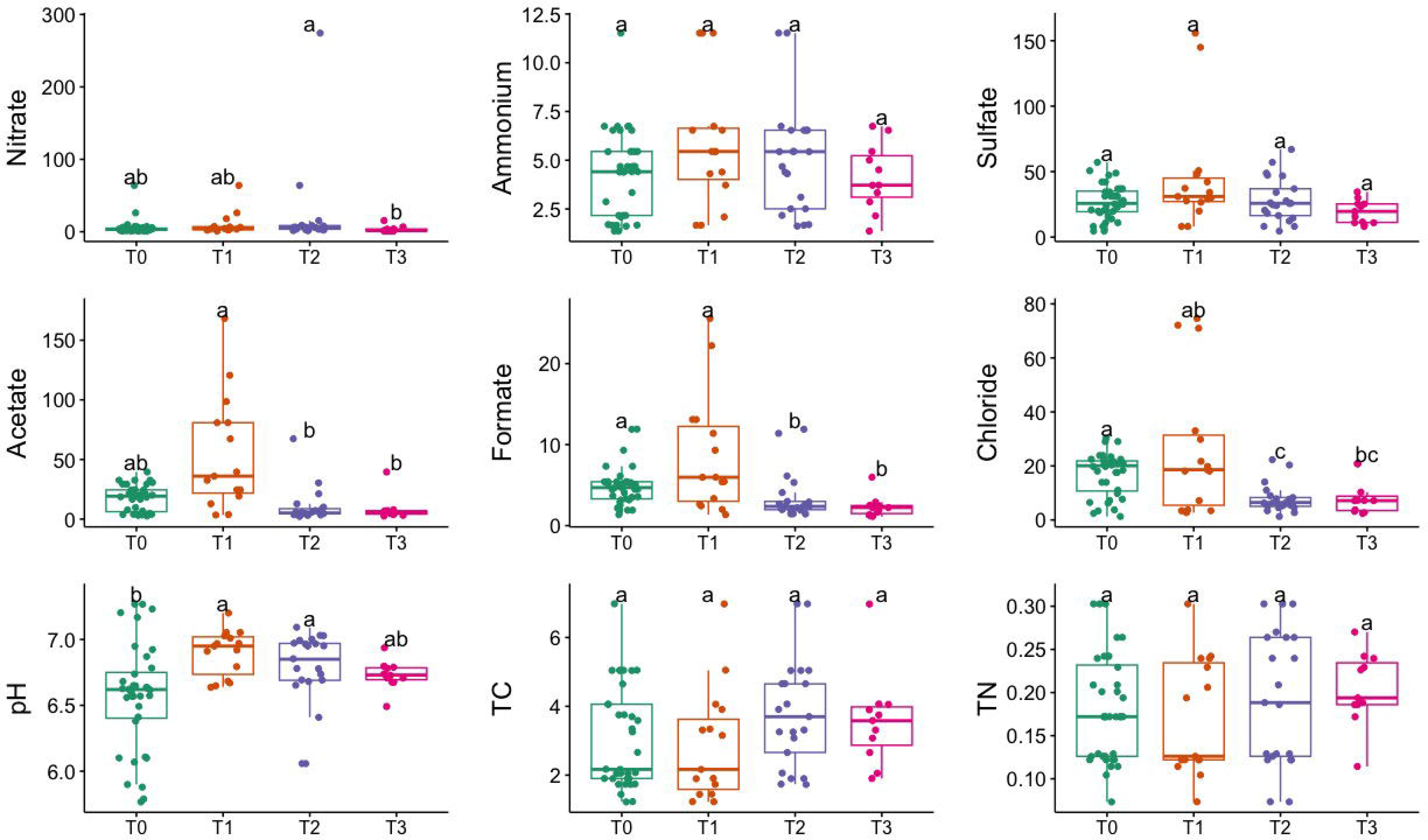
Temporal variation in soil porewater geochemistry, pH, Total Carbon (TC), and Total Nitrogen (TN) measurements. Different letters mean significant differences, and the same letters mean no significant difference (p < 0.05, KW-Dunn)

### 3.2. Microbial community richness and diversity

Our MiSeq runs yielded a total of 16.9 M sequences, with a mean of 201,376 sequences/sample and a total of 33,234 ASVs after trimming, filtering, and denoising. The Shannon-Wiener curve of all samples leveled out at a sequencing depth of 5,022 sequences/sample (Supplementary Materials, Figures S2 and S3), suggesting that the richness of the samples has been fully observed. Figure 3 shows the alpha diversity variation in the microbial community across different time points (T0 - Pre-Burn, T1-Immediately After Burn, T2 - 3 Months After Burn, T3 - 17 Months after Burn). The study revealed a significant increase in species richness from the initial (T0) to the final time point (T3). The Observed richness (ASVs) exhibited a significant rise, with counts increasing from 2051 at T0 to 2595 at T3 (p = 0.003). The Chao1 richness estimator mirrored this trend, with the estimate increasing from 2053 at T0 to 2598 at T3. The Shannon index showed variations between time points but did not reach statistical significance (p = 0.16). PERMANOVA was applied to the differential test of weighted Unifrac distances among groups and showed that there was a significant difference across time points (R^2^= 0.118, F= 3.64, p = 0.001) but less significantly so for the types of additional slash fuel (R^2^= 0.061, F= 3.64, p = 0.018). Violin plots comparing the alpha diversity index can be found in supplementary materials (Figure S4).

**Figure 3.**
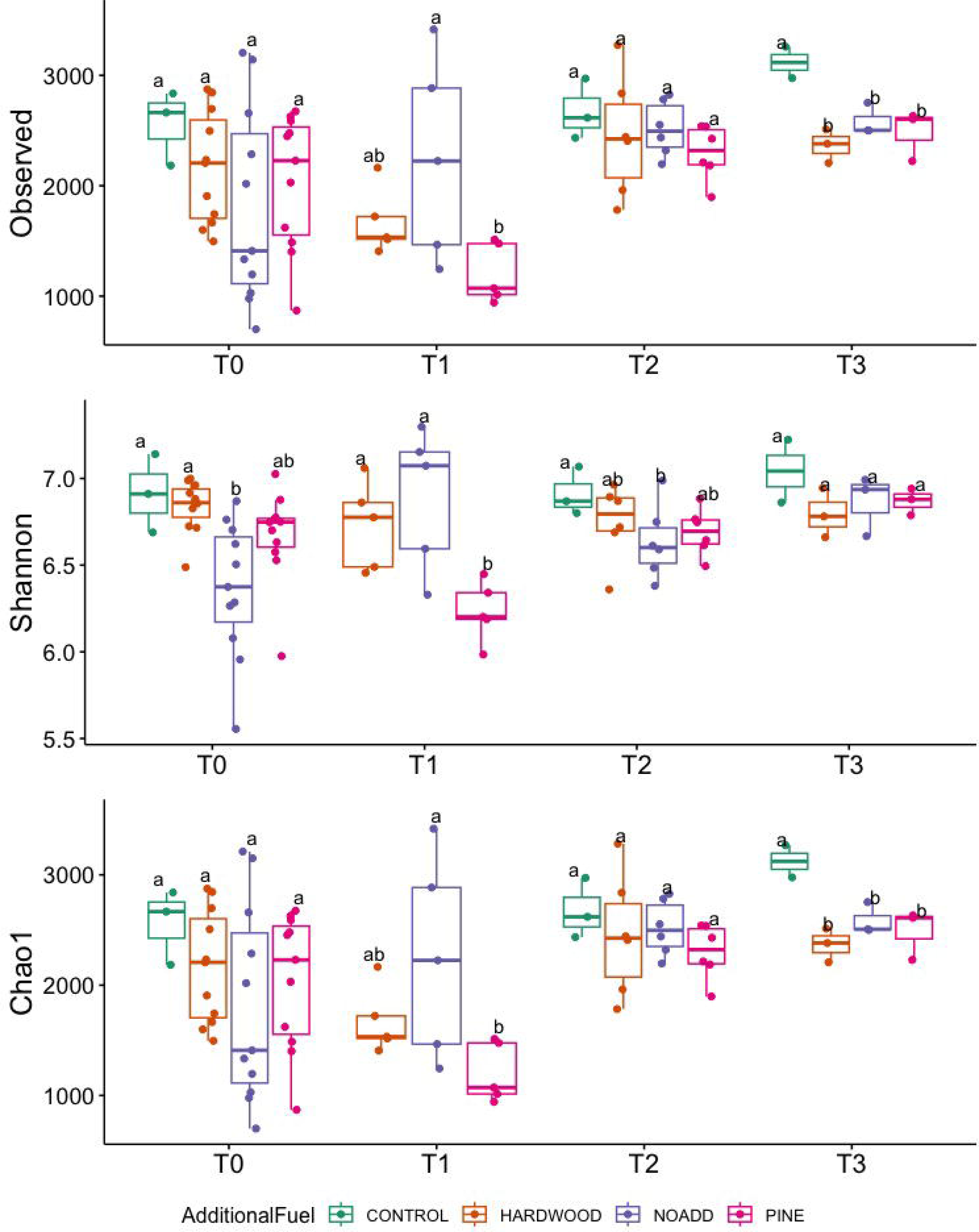
Temporal variation in Observed Richness, Shannon diversity index, and Chao1 richness estimator across different fuel treatments (T0, T1, T2, T3). Different letters mean significant differences, and the same letters mean no significant difference in abundance (based on ANOVA followed by Duncan’s multiple range test, p < 0.05).

### 3.3. Differences in bacterial community composition

Figure 4 illustrates the normalized relative abundance of bacterial phyla across the community using ANOVA to identify to any statistical significance over time. Considering the taxonomic abundance across different time points, the three most predominant phyla were Proteobacteria, Acidobacteria, and Actinobacteria, with mean relative abundances of 29%, 25%, and 13%, respectively. Differential testing showed distinct community differences across groups (alternate relative abundance plots utilizing a different method can be found in Figures S6 A – D). Small but significant increases were observed between post-fire time points for Proteobacteria (T0 – 28.96%, T2 – 31.64%) and Acidobacteria (T0 – 22.04%, T3 – 28.04%) (p < 0.05, ANOVA). The relative abundance of the bacterial phylum Verrucomicrobia decreased from 8.21% at T0 to 5.55% % at T3 (p < 0.05, ANOVA). Taxonomic bar plot without application of differential tests can be found in Supplementary Materials, Figure S5.

**Figure 4.**
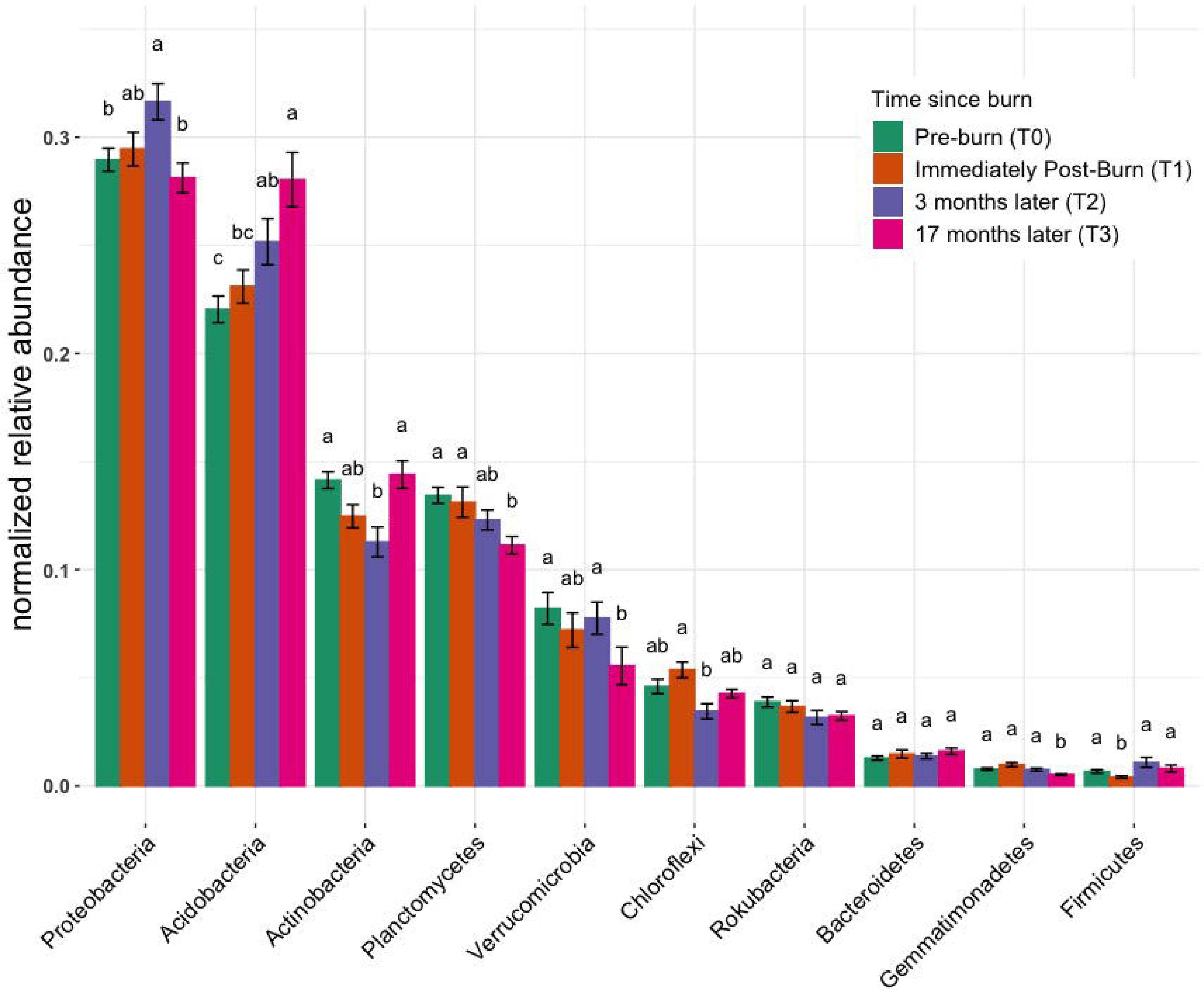
Normalized relative abundance across samples. These bar charts provide insights into the distribution of taxonomic groups among samples, elucidating shifts in microbial community composition. Different letters mean significant differences, and the same letters mean no significant difference (based on ANOVA followed by Duncan’s multiple range test, p < 0.05).

### 3.4. Relationships between microbial community composition, function, and soil chemical properties

Microbial communities at different time points all clustered very closely together in the NMDS plot (Figure 5), suggesting little to no differences in pre- and post-fire microbial communities. The NMDS ordination, RDA, and correlation analyses revealed significant differences between specific bacterial phyla (illustrated in Figure 5 and Figure 6) and soil properties (PCoA ordination was also used to visualize the weighted Unifrac matrix. Please see Figure S7 in Supplementary Materials for the ordination plot). The RDA triplot (R^2^ = 0.506, adjusted R^2^ = 0.302) in Figure 6 explains 30% of the community composition regarding species abundance across samples. Looking at similarities between the time points within the response matrix, we notice much overlap between time points across samples, which suggests that the communities have not diverged with time since the burn. Also, looking at the shaded ellipses for different time points, the clusters are noticeably closer together with more time since the burn, with the only exception being the extreme situation immediately after the burn. Phyla that are closer together, such as Acidobacteria and Bacteroidetes, compared to Proteobacteria and Actinobacteria, occupy more sites in common. Longer arrows are noted for environmental variables such as pH, Formate, Acetate, and Ammonium, indicating that these variables strongly drive the variation within the community matrix. For the correlation analysis, after carrying out a Mantel test, only the significant correlations are elaborated upon here. Soil pH was positively correlated with the Acidobacteria, Azohydromonas, and Chloroflexi phyla while exhibiting a negative correlation with the Acidothermus, Acidipila, Mycobacterium, and Bradyrhizobium genera. Na^+^ concentrations were significantly positively correlated to Azohydromonas, Solimonas, and Raoultibacter. Furthermore, porewater NO_3_^−^ concentrations positively correlated with the Verrumicrobium, Anaerocolumna, and Methanosaeta genera. In contrast, porewater POL³L concentrations exhibited positive correlations with various genera, including Ferrovibrio, Labilithrix, and Uliginosibacterium (Correlation Heat map can be found in Supplementary Materials, Figure S8). No significant differences were observed in terms of correlation between Total C and N concentrations concerning different taxa.

**Figure 5.**
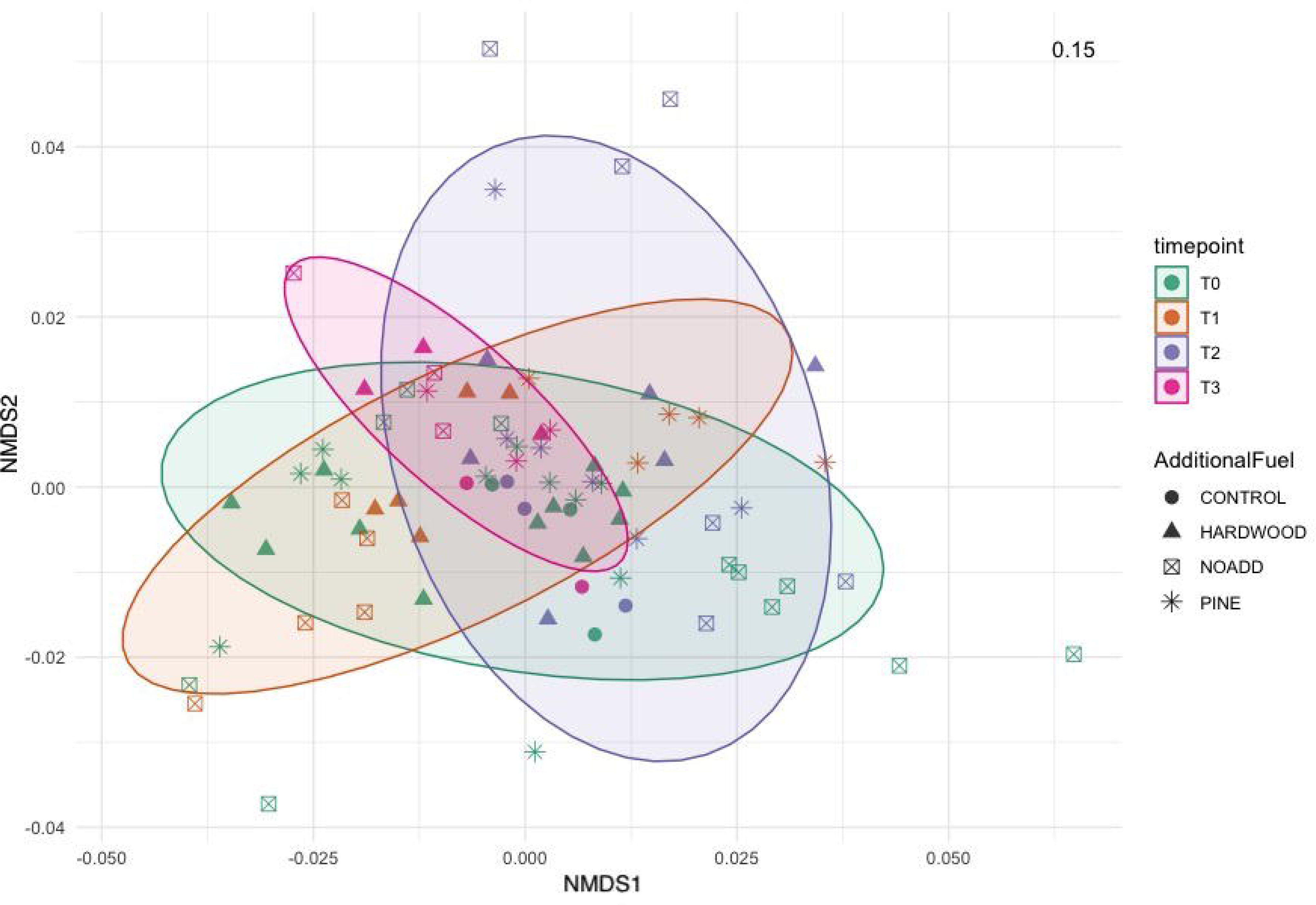
Non-metric Multidimensional Scaling (NMDS) ordination based on weighted Unifrac distance, showing the bacterial community structures derived from relative abundance based on phylum level across the different samples. The stress value denotes the goodness of fit.

**Figure 6.**
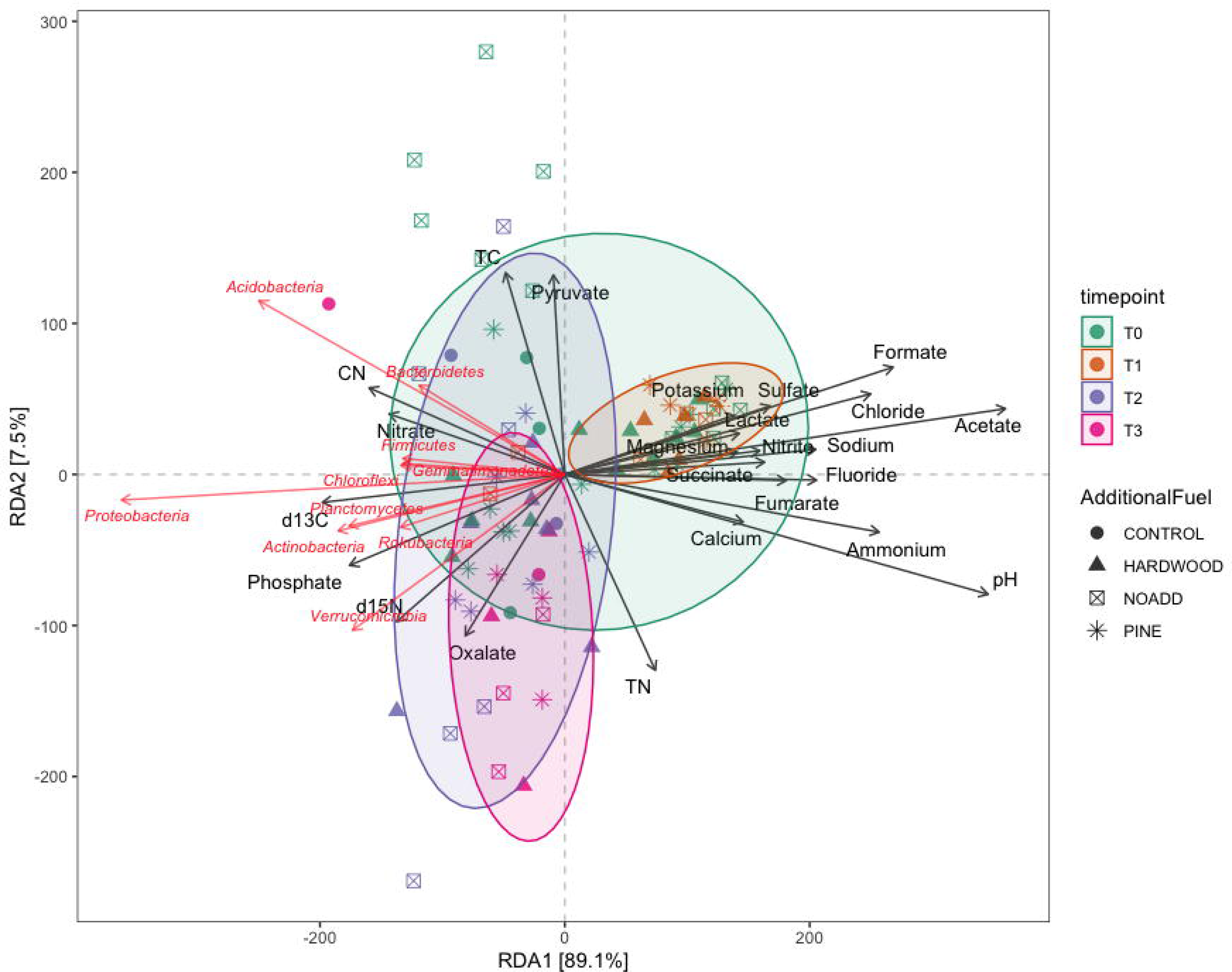
Redundancy analysis (Bolyen et al.) triplot illustrating multivariate associations between environmental factors and microbial composition. Dots symbolize samples, while black arrows depict environmental parameters, and red arrows denote microbial families, with the arrows indicating the correlation between environmental variables and distinct bacterial families.

To investigate potential biotic nutrient cycling mechanisms occurring in our samples, we used the FAPROTAX database to match genomic functions with the taxonomy derived from our 16S amplicons. Functional traits with the ten highest relative abundance of ASVs have been shown in Figure 7. These results suggest dynamic shifts in the abundance levels of aerobic ammonia oxidation traits following the burn event. There is a significant enrichment in the abundance of these traits immediately after the burn (p < 0.05, ANOVA). However, over time, the abundance levels of aerobic ammonia oxidation traits exhibit a notable decline, falling below the initial relative abundance levels at subsequent time points, at 3 months and 17 months post-burn. Similar trends were observed for nitrification traits, indicating a broader impact on nitrogen cycling processes in the soil following the prescribed burn. For more visualizations of putative functional traits and correlation with environmental variables, see Figure S8 of Supplementary Materials.

**Figure 7.**
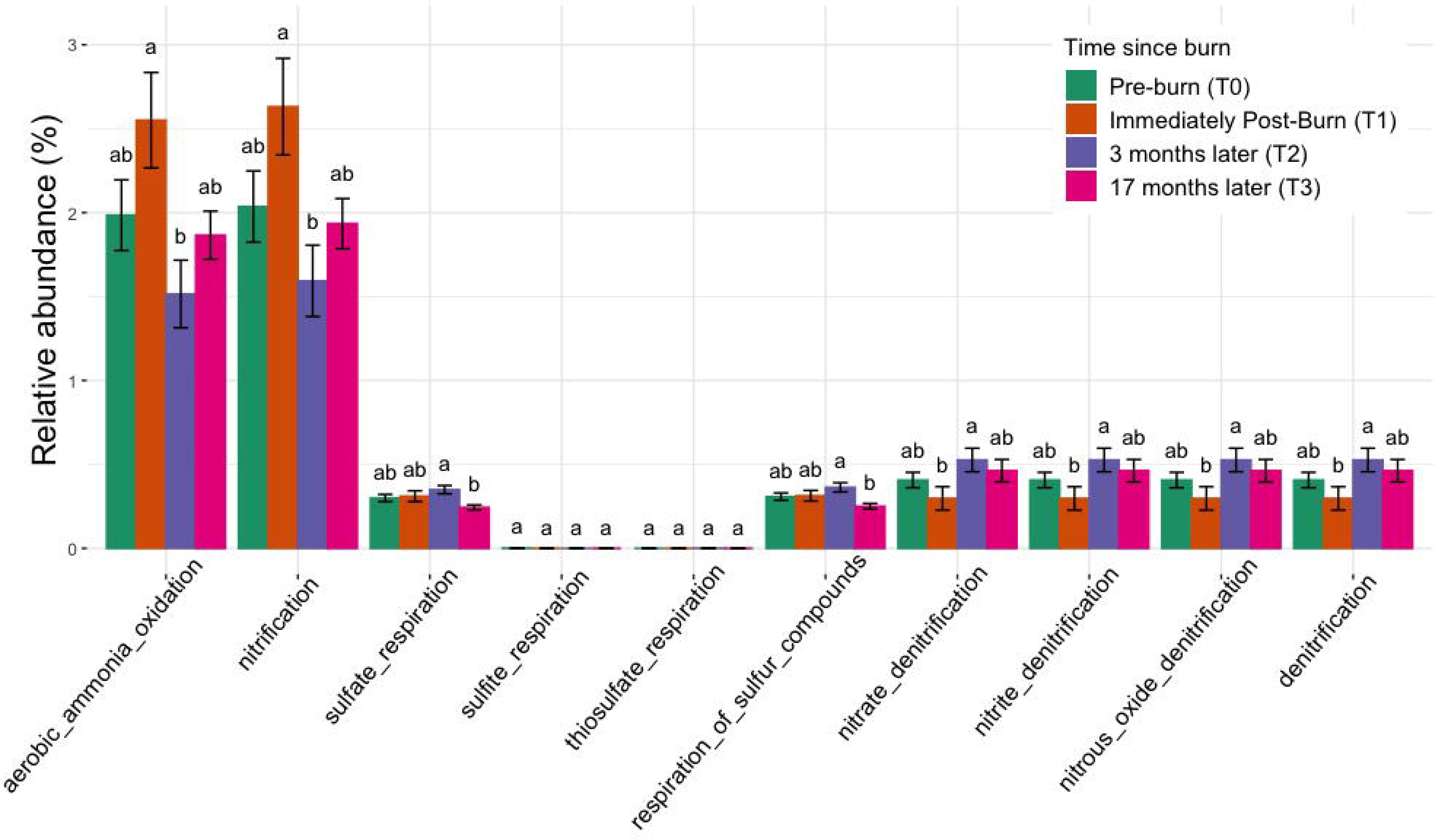
Normalized relative abundance bar plot using FAPROTAX shows the enrichment and depletion of specific functional traits across all taxonomic groups over time. Significant changes were observed across time points for nitrogen cycling and sulfate respiration in post-burn soils (based on ANOVA followed by Duncan’s multiple range test, p < 0.05). Different letters mean significant differences, and the same letters mean no significant difference.

## 4. Discussion

We demonstrated that low-intensity fire has a significant impact on both soil chemistry and the soil microbial community. Porewater concentrations for soluble ions exhibited notable stability, with minor variations discernible across different fuel types and time points. Nevertheless, significant changes were observed in Acetate, Formate, and Nitrate concentrations, suggesting discernible modifications in the solute chemistry of the soil post-fire. These changes in solute chemistry may have been caused by the release of organic compounds during the burn, leading to increased acetate levels (Certini, 2005). The presence of NO_3_^−^ in higher concentrations could be attributed to the heat-induced transformation of soil organic matter, resulting in the release of these ions into the porewater (Hubbert et al., 2015). An immediate increase in NH_4_^+^ concentration and the subsequent return to baseline levels over an extended time was reported in multiple different studies (Knoepp and Swank, 1993; Klimas et al., 2020). In the case of low-intensity burns, the organic nitrogen fraction tends to be more affected than the inorganic fraction, as has been reported by multiple studies, and our results seemingly concur (Wan et al., 2001; Taylor and Midgley, 2018). NH_4_^+^ and NO_3_^−^ concentrations in soil usually increase following a fire event due to the incomplete combustion of nitrogenous materials, including vegetation and denatured soil proteins (Certini, 2005; Johnson et al., 2011; Fontúrbel et al., 2012). This corroborates our study’s findings, where we noticed similar statistically significant changes in NO_3_^−^ porewater concentration post-burn. It is imperative to note here that in the results of the prediction of putative microbial functions using FAPROTAX, there is significant enrichment in the relative abundance of nitrification and aerobic ammonia oxidation traits, and these implications are further discussed later in this section.

Changes in soil porewater concentration of the base cations in this study can be attributed to ash incorporation soil or location-specific characteristics (Neary et al., 1999; Certini, 2005; Taylor and Midgley, 2018). The total carbon content showed an initial increase post-burn for Pine Slash and No Added Slash treatments, followed by a decline at the 17-month mark, indicating dynamic changes in organic carbon turnover in response to fire. The modification of soil carbon content in response to wildfires can be attributed to introducing of novel carbon sources, notably charred organic matter and plant debris (Certini, 2005) and observed oscillations in total carbon content may allude to dynamic shifts in the equilibrium between carbon inputs, including plant-derived materials and organic substrates, and carbon outputs, which predominantly manifest through microbial decomposition processes within the post-fire landscape.

Our observations reveal shifts in alpha diversity following prescribed fires, reflecting fluctuations in the Shannon index, Chao1 estimator, and observed richness. These findings align with research suggesting microbial community structure and diversity alterations following fire events (Balser and Firestone, 2005). While Shannon index variations were not statistically significant, the Chao1 estimator and observed richness suggested an increase in potential taxonomic richness over time, contributing to functional diversity. Although the Shannon index showed variations between time points, which implies dynamic changes in the composition and structure of the microbial community following the burn event, the lack of statistical significance suggests that there might not be significant changes in evenness across the different stages of ecological succession. The Chao1 estimator mirrored the trend observed in observed richness, implying an increase in potential taxonomic richness over time and implications for the functional diversity of soil bacterial communities (Tripathi et al., 2018). The trends in observed richness support this notion, with an overall increase in richness by the final timepoint, and there was a noticeable drop right after the burn event, which can be explained by the physical effects of the fire event. These results align with previous studies that have reported increases in microbial diversity following fire disturbances over a medium-term time scale (Staddon et al., 1998; Fontúrbel et al., 2012).

In the context of enriched Chao1, Shannon, and observed richness over time, both stochastic and deterministic processes influence microbial community dynamics. While stochastic processes may contribute to initial community assembly, the sustained increase in richness suggests deterministic factors such as niche differentiation and biotic interactions play a pivotal role in shaping the microbial community structure over time (Dini-Andreote et al., 2015). Enriching microbial diversity over time implies that deterministic processes such as niche differentiation, environmental filtering, and biotic interactions likely influence community structure. Deterministic processes may drive the observed patterns of increased richness and diversity, reflecting adaptations to changing environmental conditions and interspecies interactions. Prescribed fires could enhance functional redundancy within soil bacterial communities, promoting ecosystem stability and resilience, as suggested by prior investigators (Delgado-Baquerizo et al., 2018). Increased alpha diversity suggests the introduction of additional microbial taxa with similar functional traits, which ensures the continuity of essential ecosystem functions in a post-disturbance environment. This redundancy enhances the stability and resilience of ecosystem processes, such as organic matter decomposition and nutrient cycling (Allison and Martiny, 2008; Philippot et al., 2010).

Furthermore, shifts in alpha diversity following prescribed fires may lead to more diverse functional traits within soil bacterial communities. This diversification broadens the range of ecological functions, including the breakdown of complex organic matter, nitrogen fixation, and disease suppression (Schimel and Schaeffer, 2012; Louca et al., 2017). Changes in functional traits will be further discussed later when we analyze the enrichment of putative microbial functions using the FAPROTAX database. The impact of prescribed fires extends beyond mere diversity shifts, playing a crucial role in shaping the functional dynamics of soil ecosystems.

The taxonomic composition analysis reveals that Proteobacteria, Acidobacteria, and Actinobacteria were the dominant phyla in the microbial communities across all time points. Furthermore, differential tests indicate significant shifts in the relative abundance of specific phyla. Proteobacteria and Acidobacteria showed small but significant increases in relative abundance between pre-fire and post-fire time points, suggesting their potential roles in post-fire soil recovery through natural succession-driven processes (Jones et al., 2009; Moyano et al., 2013). In contrast, Verrucomicrobia exhibited a significant decrease in relative abundance over time. This decline may be related to their sensitivity to environmental changes, as Verrucomicrobia are known to be associated with more stable and undisturbed soil conditions (Bergmann et al., 2011). These findings are consistent with previous studies that have reported shifts in the relative abundances of these phyla in response to fire disturbances (Lauber et al., 2008; Weber et al., 2014).

Correlation analyses revealed significant associations between specific bacterial phyla and soil properties. Soil pH, a fundamental soil characteristic, exhibited positive correlations with Acidobacteria and Chloroflexi, emphasizing their potential preference for specific pH ranges (Fierer and Jackson, 2006; Aponte et al., 2022). Conversely, Acidothermus, Acidipila, Mycobacterium, and Bradyrhizobium genera displayed negative correlations with soil pH, suggesting their sensitivity to pH fluctuations. These correlations highlight the complex interplay between microbial community composition and soil pH (Lauber et al., 2008). Porewater NO_3_^−^ concentrations positively correlated with Verrucomicrobium, suggesting their potential involvement in nitrogen cycling in post-fire soils (Bergmann et al., 2011; Mohammadi et al., 2017).

In contrast, porewater POL³L concentrations positively correlated with genera such as Ferrovibrio and Uliginosibacterium, indicating their potential roles in phosphorus dynamics. These correlations underscore the importance of specific microbial taxa in mediating nutrient cycling processes in response to prescribed burns (Cederlund et al., 2014). The changes in grouped relative abundance for the functional traits predicted using the FAPROTAX database indicate a potential response to the disturbance caused by the fire. This enrichment may reflect heightened microbial activity or proliferation of bacteria involved in aerobic ammonia oxidation processes in response to post-burn conditions. The subsequent depletion of the traits is interesting as this downward trend suggests a gradual depletion or decline in the populations of microorganisms associated with aerobic ammonia oxidation traits as the soil ecosystem undergoes post-disturbance recovery and stabilization.

Our PERMANOVA analysis revealed significant differences in microbial community structure across the four-time points. The variation observed in community structure emphasizes the dynamic nature of soil microbial communities following prescribed burns. However, it is noteworthy that the additional slash fuel had a limited impact on microbial communities, suggesting that other factors, such as fire severity or soil properties, might exert stronger influences on post-burn microbial community structure. Furthermore, the clustering of microbial communities in NMDS ordination and RDA analysis (Figure 5 and Figure 6) at different time points indicates that the overall pre- and post-fire microbial communities remained relatively consistent (For PCoA ordination plot, please see supplementary material, Figure S4). This resilience of soil microbial communities may be attributed to factors such as resilient microbial taxa or the rapid post-fire recolonization of microbes (Whitman et al., 2016; Whitman et al., 2019). Specific taxa did respond to fire-induced changes in soil properties, but the overall structure and function of the microbial community remain relatively consistent. Such stability could be attributed to microbial populations’ resilience and ability to adapt to changing conditions. It is important to note here that the longer vector arrows for pH and Acetate indicate that these two explanatory variables are strong drivers of microbial community composition in temperate mixed forest soils, with pH being the highest determinant factor explaining the most variations in the community composition. The relative abundance of some of the phyla, like Bacteroidetes, Acidobacteria, and Firmicutes, were correlated with increasing NO_3_^-^ concentrations. (Weber et al., 2014) mention that variations in the abundance of Bacteroidetes can be partially explained and correlated to carbon mineralization rates. An increase in abundance was observed for Bacteroidetes with each subsequent time point, albeit these changes were minor. Looking at the RDA triplot, we notice that it is correlated to the carbon-nitrogen (CN) ratio. This observation agrees with the carbon cycling characteristics expected of an early-stage post-fire soil ecosystem, as noted by (Fierer et al., 2007; Smith et al., 2008) since microbial anabolism is facilitated by the release of labile carbon into the soil after a burn event.

## 5. Conclusion

Our study of the impacts of slash fuel type and time on prescribed fire in Southern Appalachian Forest soils echoes the findings of similar studies. The prescribed burns triggered transient changes in specific soil properties and microbial communities, which largely reverted to their pre-fire conditions over a 17-month observation period. Bacterial communities and composition exhibited resilience to these prescribed burns, but noteworthy alterations were observed in the relative abundance of specific microbial taxa. These results substantiate the prevailing view that prescribed fires have limited effects on soil microbial community diversity and composition, especially within ecosystems well-adapted to fire. While our study has provided valuable insights into the impacts of time since and slash-fuel-type of prescribed fire on temperate mixed forests, it is essential to acknowledge that its effects on soil properties and microbial communities are complex and multifaceted. This research adds to the growing body of knowledge on the subject, emphasizing the resilience of these ecosystems to low-intensity prescribed burns. However, to fully grasp the long-term implications and sustainability of prescribed fire as a forest management tool, further investigation into the extended impacts of repeated burning events on nutrient cycling and ecosystem dynamics in temperate mixed forests is imperative. Such studies will aid in developing informed land management strategies that balance ecological conservation with the need for forest health and resilience in the face of evolving environmental challenges.

## Acknowledgements

The authors thank the Office of Research at the University of Tennessee, Knoxville, for funding the project via a FUSION seed grant. We would also like to thank the Institute for a Sustainable & Secure Environment for funding extension work on this project.

