## Supplementary Document for "Impact of Prescribed Fire on Soil Microbial Communities in a Southern Appalachian Forest Clearcut"

Supplementary Material

### 1 Supplementary Figures and Tables

#### 1.1 Supplementary Figures

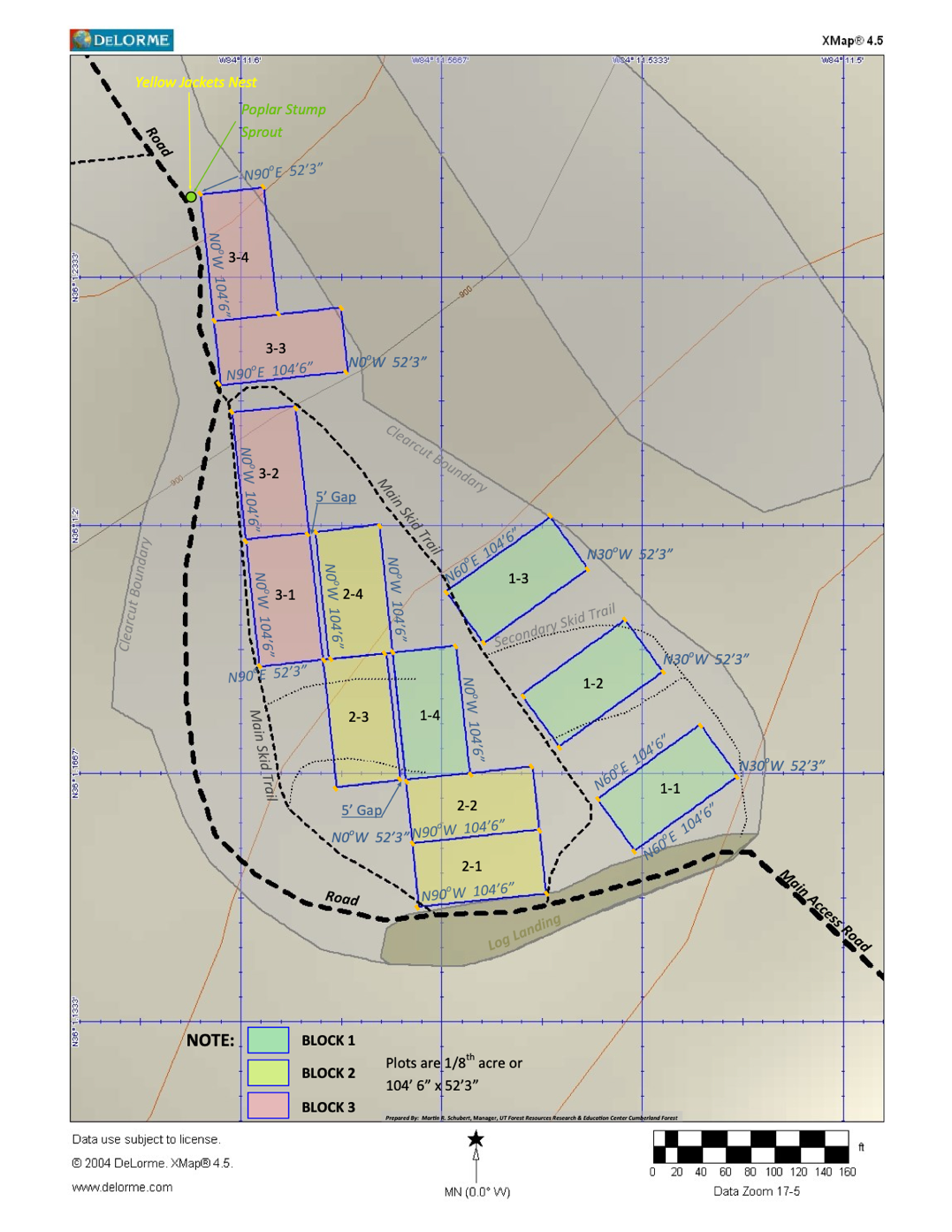

Figure S 1
Detailed Site map for FRREC Burn Site showing plot arrangement and size details. Plots 1-1, 2-3 and 3-2 had additional hardwood slash, Plots 1-2, 2-1, and 3-1 used pine slash, Plots 1-3, 2-4, and 3-4 had no additional slash fuel while plots 1-4, 2-2 and 3-3 were control plots.

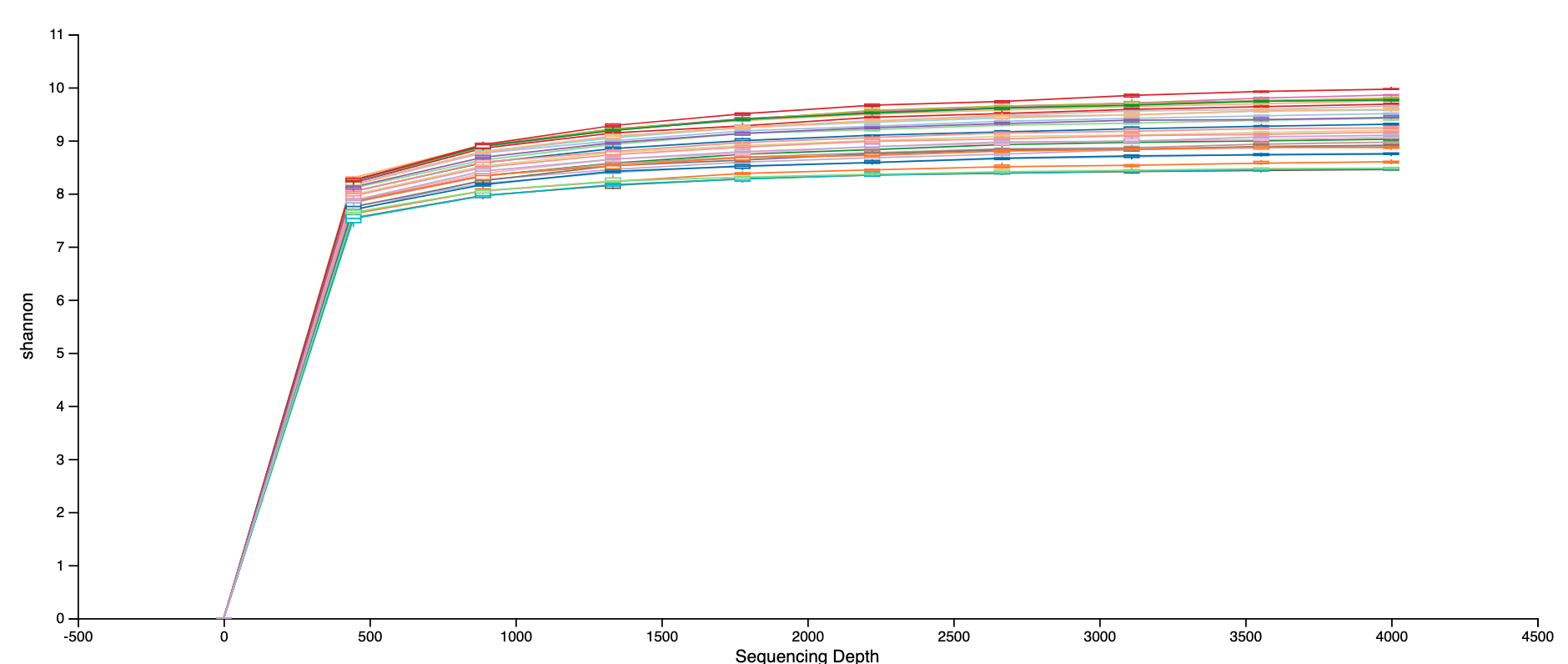

Figure S 2
Alpha Rarefaction Plot for First 16S Sequencing Run carried out in 2018. All samples have been represented with individual colors, and all curves eventually levelled out at sequencing depth shown.

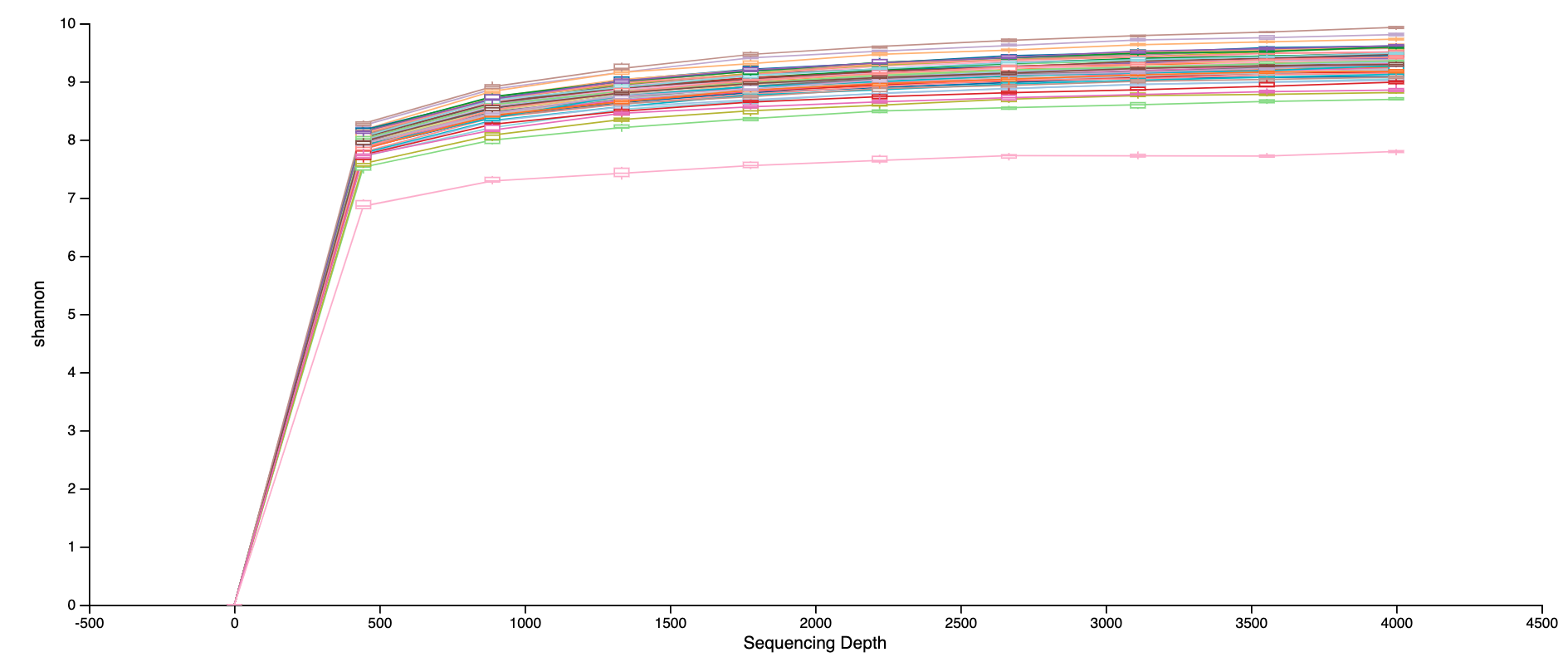

Figure S 3

Alpha Rarefaction Plot for First 16S Sequencing Run carried out in 2019. All samples have been represented with individual colors, and all curves eventually levelled out at sequencing depth shown.

**Figure S 4**

Violin plots illustrating fluctuations in alpha diversity indices (ACE, Chao1, Observed, Pielou, Shannon, Simpson) across distinct slash fuel types and timepoints. Statistical significance was assessed using Wilcoxon test.

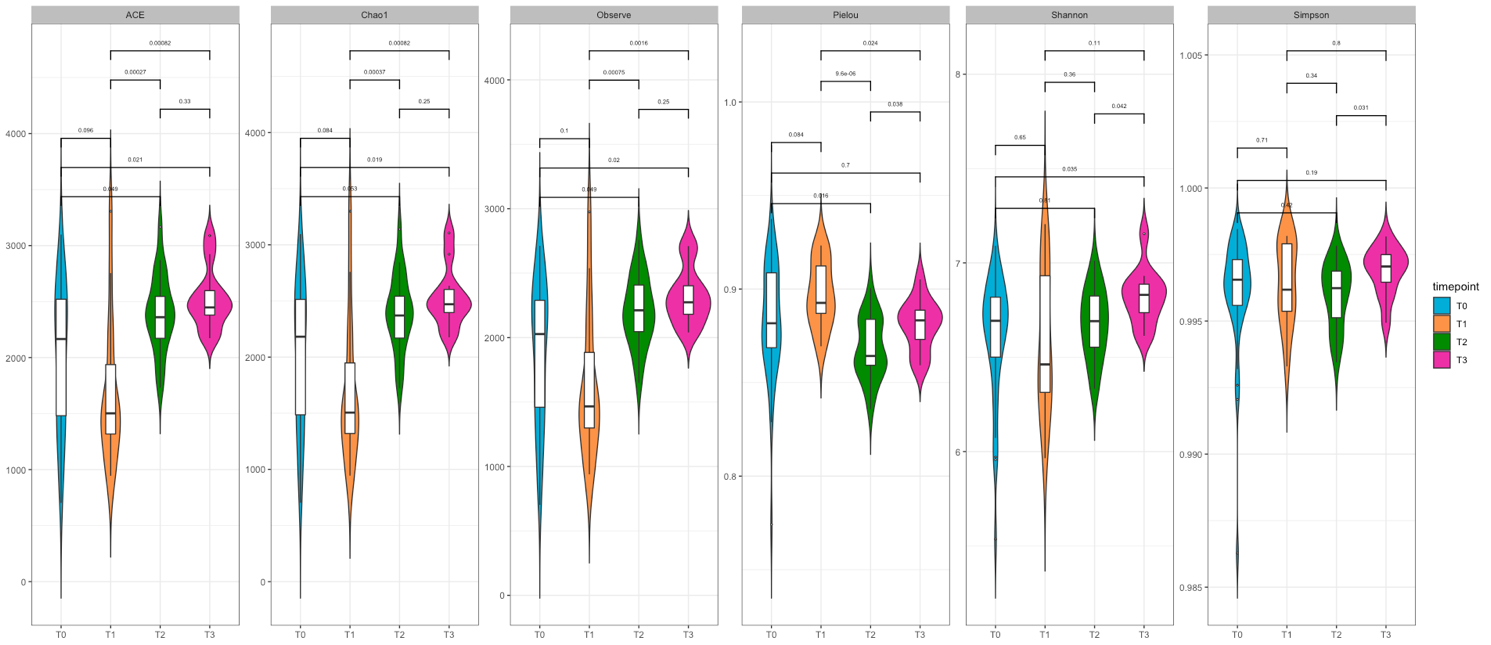

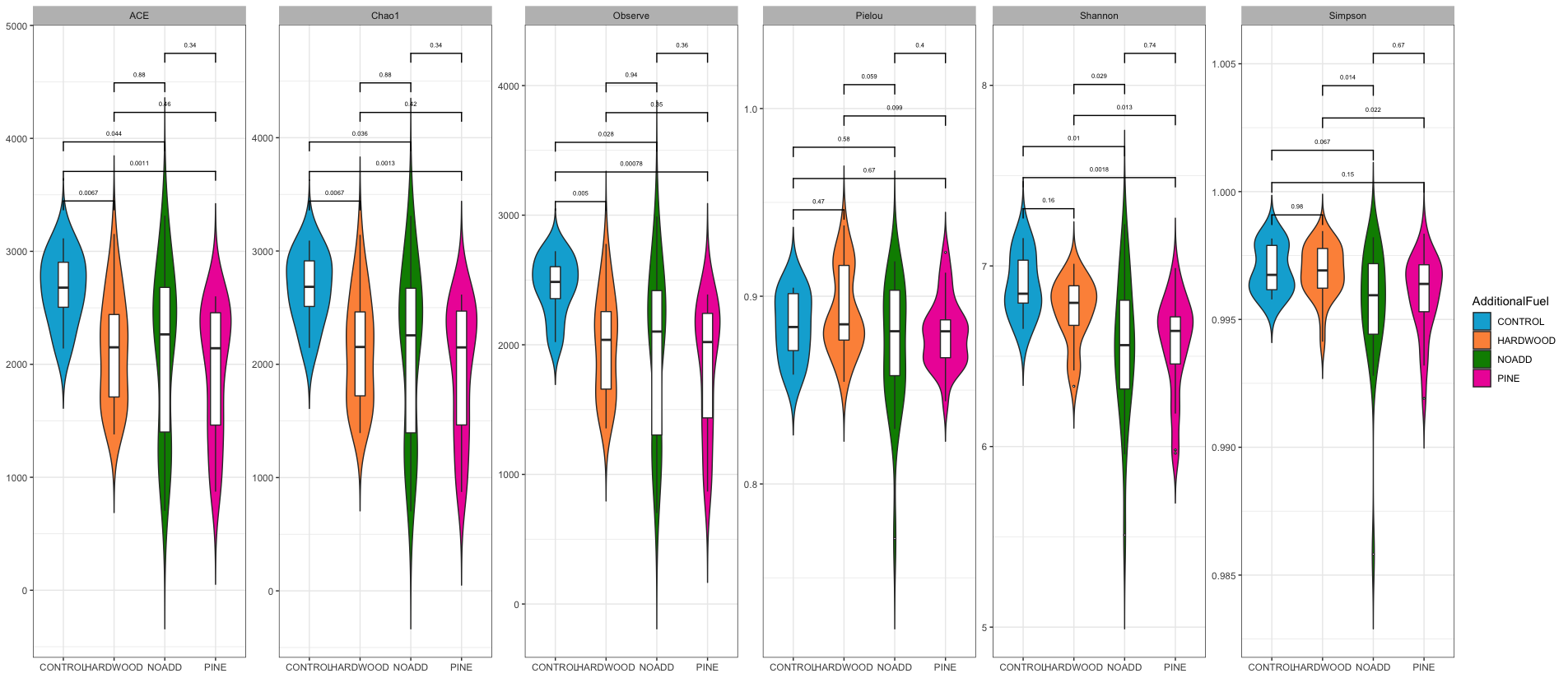

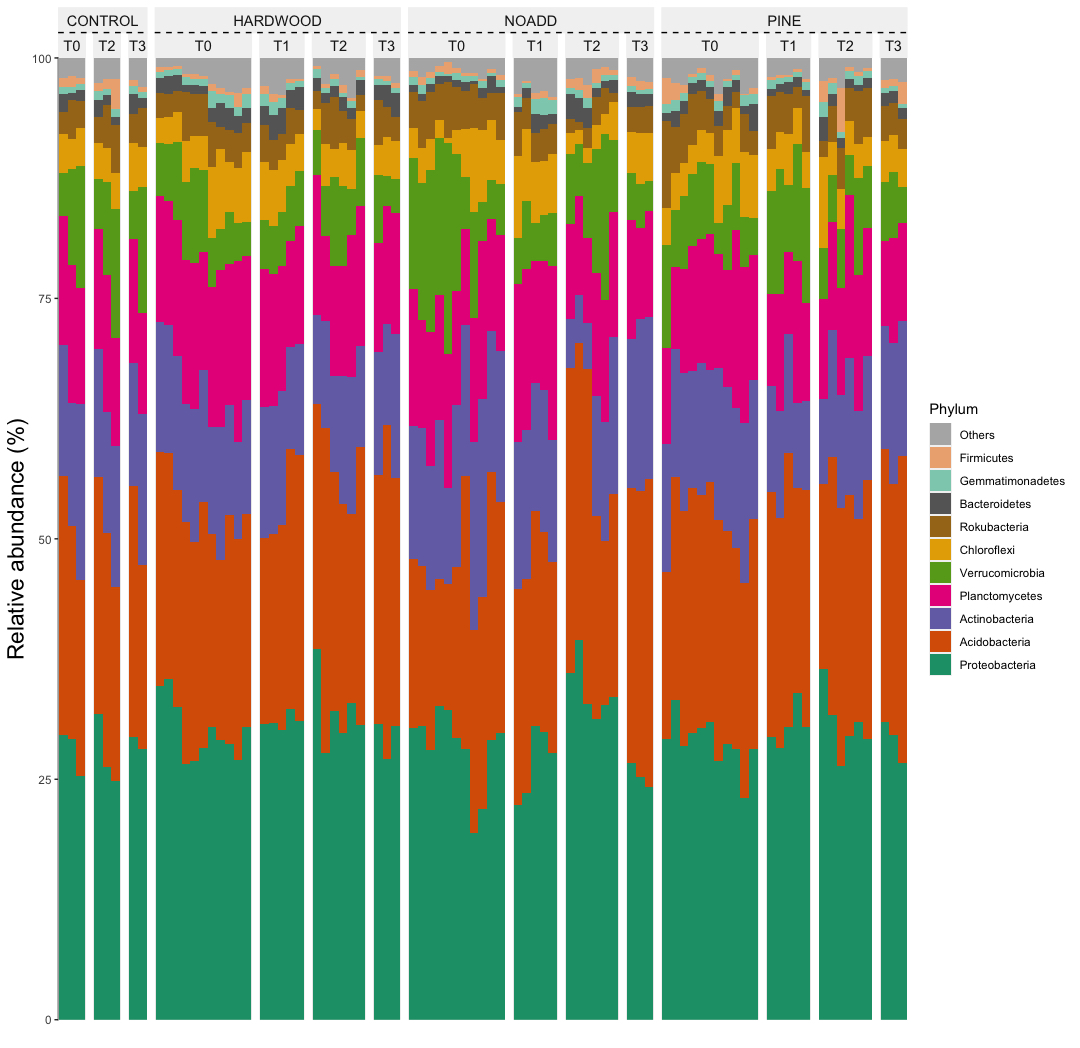

Figure S 5 Taxonomic Abundance chart. The top 10 phyla were selected to provide insights into the distribution of taxonomic groups among samples as arranged by timepoints and additional fuel treatment type.

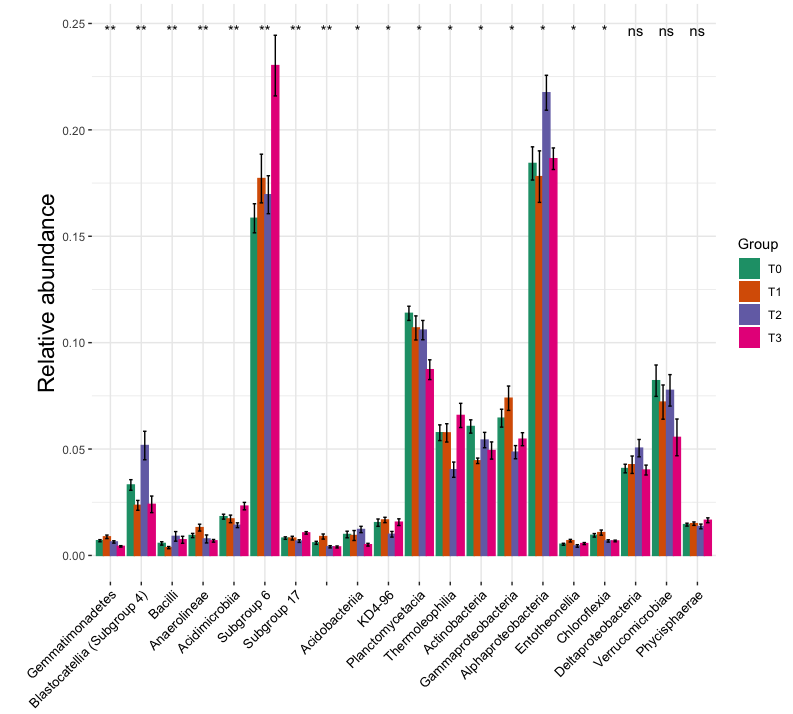

Figure S6 A

Relative Abundance plot at Class level, generated using output from the ALDEx2.kw method in the microeco package. The method performs a Kruskal-Wallis test and a glm ANOVA for the differential test metrics. Asterisks denote significant difference from the control, *P < 0.05; ** P < 0.01.

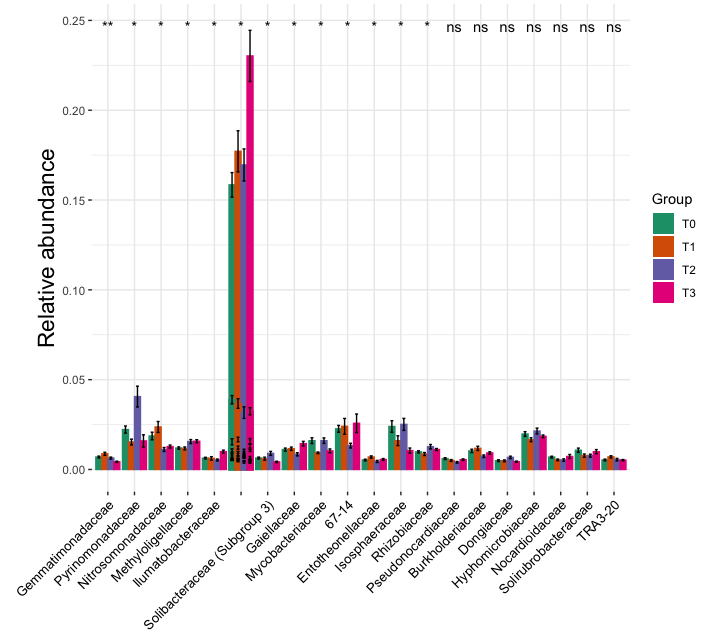

Figure S6 B

Relative Abundance plot at Family level, generated using output from the ALDEx2.kw method in the microeco package. The method performs a Kruskal-Wallis test and a glm ANOVA for the differential test metrics. Asterisks denote significant difference from the control, *P < 0.05; ** P < 0.01. Non-significant results are denoted by “ns”.

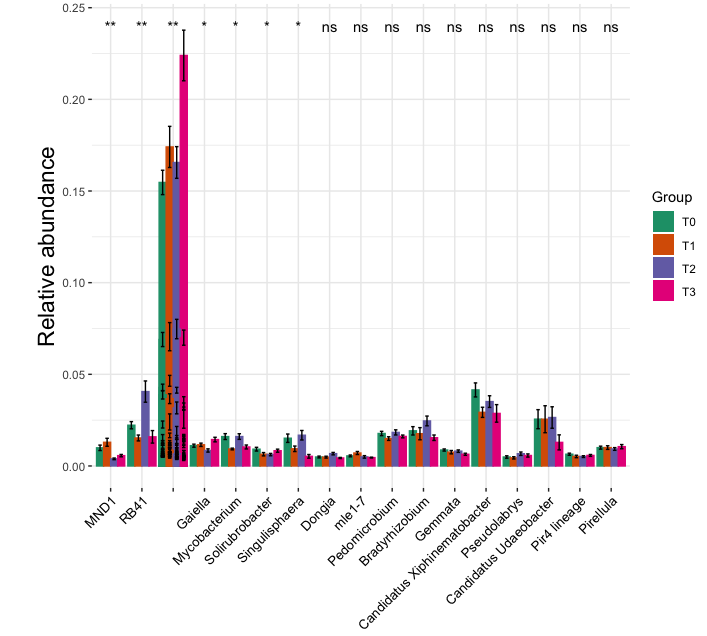

Figure S6 C

Relative Abundance plot at Genus level, generated using output from the ALDEx2.kw method in the microeco package. The method performs a Kruskal-Wallis test and a glm ANOVA for the differential test metrics. Asterisks denote significant difference from the control, *P < 0.05; ** P < 0.01. Non-significant results are denoted by “ns”.

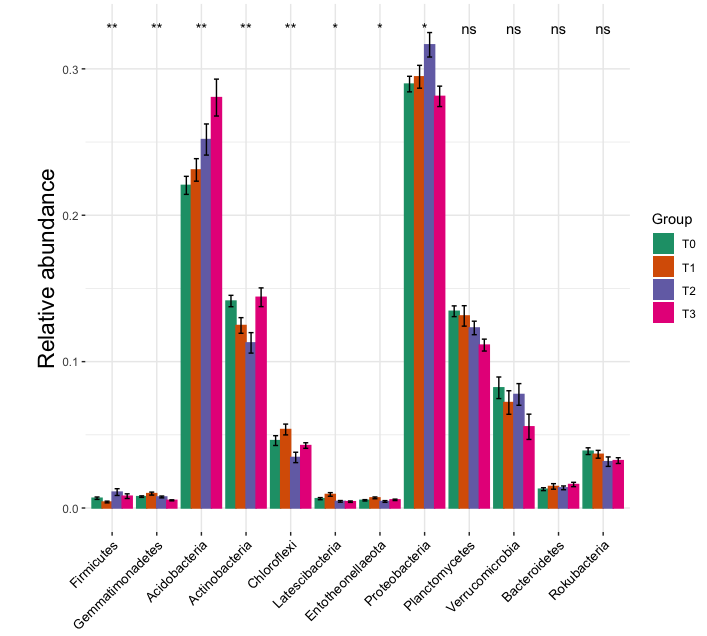

Figure S6 D

Relative Abundance plot at Phylum level, generated using output from the ALDEx2.kw method in the microeco package. The method performs a Kruskal-Wallis test and a glm ANOVA for the differential test metrics. Asterisks denote significant difference from the control, *P < 0.05; ** P < 0.01. Non-significant results are denoted by “ns”.

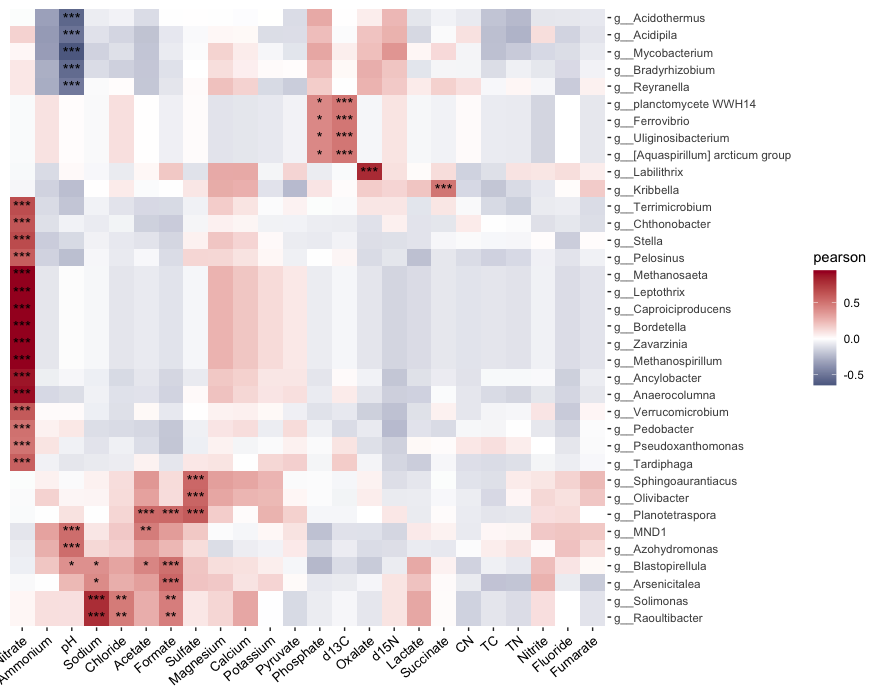

Figure S 7

Correlation heatmap depicting relationship between diverse genera and environmental parameters. Positive associations are shown in deeper red hues and negative relationships in deeper blue shades. Asterisks denote significant difference from the control, *P < 0.05; ** P < 0.01; *** P<0.001.

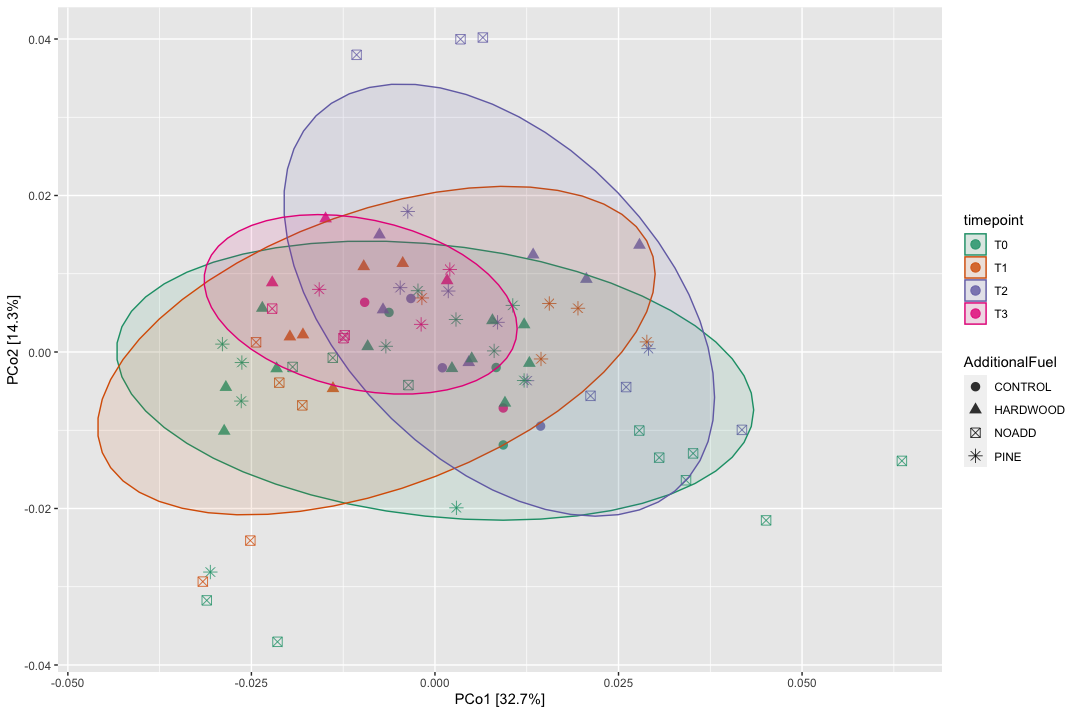

Figure S 8
Beta diversity for prokaryotic communities from soil samples. Principal coordinate analysis (PCoA) based on weighted Unifrac distances showed that time elapsed since fire explained 32.7% of variations in the soil bacterial/archaeal communities and the type of slash fuel used explained only 14.3 % of the variation. Ellipses represent standard deviation of axis scores from timepoint centroids.

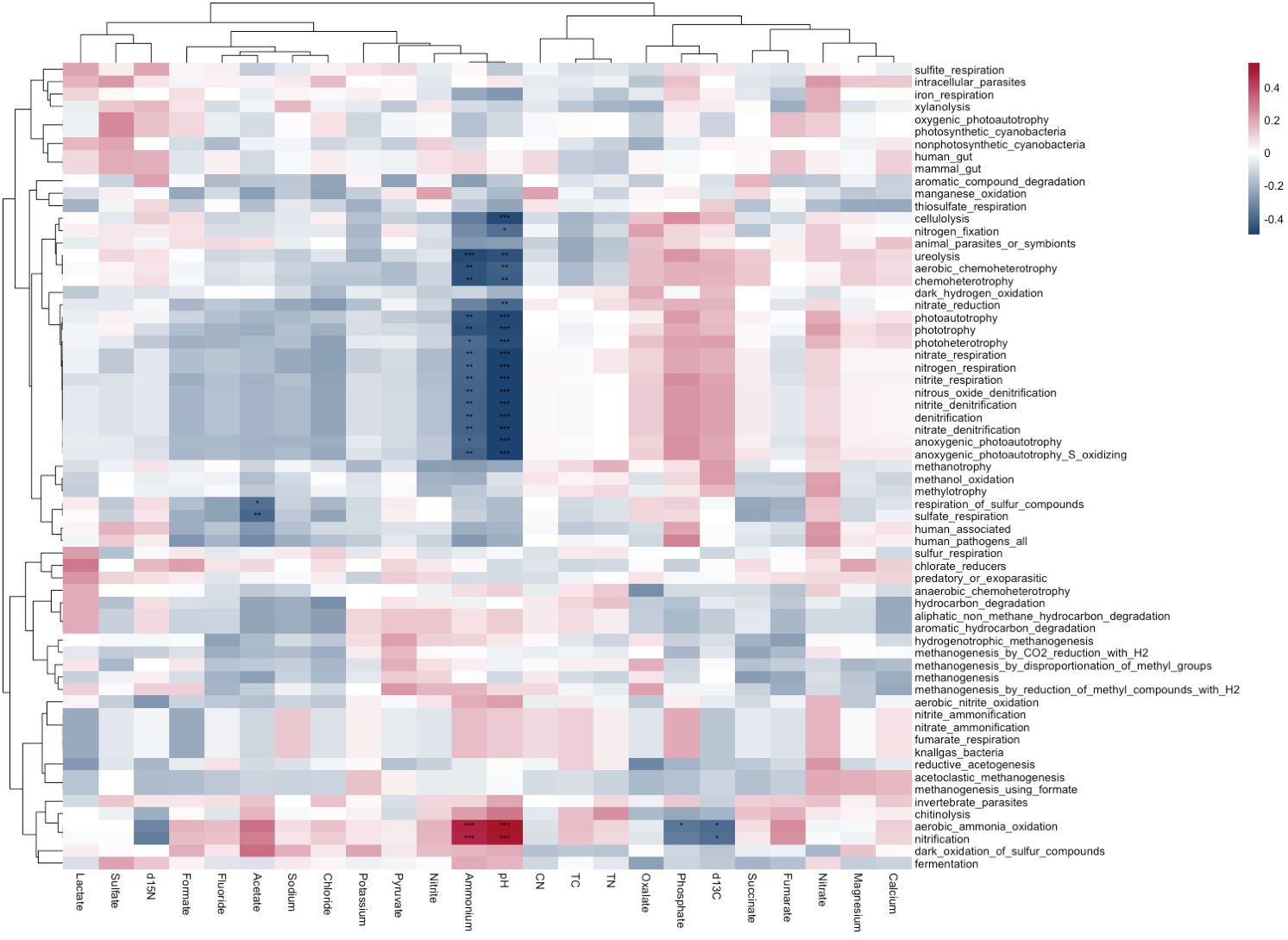

Figure S 9
FAPROTAX heat map depicting statistically significant environmental variables and their association with predicted microbial functions. The number of asterisks (*) denoting varying levels of confidence—ranging from the highest (***, p < 0.001, ANOVA) to the lowest (*, p < 0.05, ANOVA) with the color gradient indicating pearson correlation coefficient values with negative values as blue and positive values as red.

**Functional Profile of Microbial Communities with 16S dataset**

Significant correlations were noticed between acetate concentrations and sulfate/sulfur respiration (p < 0.05). As prior investigations have shown that acetate is a key substrate for sulfate respiration, it would appear. pH and NH_4_^+^ also exhibited a significant correlation (p < 0.001) with nitrogen cycling processes. This is quite understandable as pH determines the relative amounts of ammonium and ammonia ions present in the subsurface media. Fluctuations in the pH can affect the availability of these ions.

#### Supplementary Tables

Table T 1 showing the output of the cal_diff function in microeco package used to test the significance changes in environmental variables across groups. Significant differences are indicated by different letters (p<0.05, KW-Dunn) and are showed in Table T2, with the calculated P values shown here

| Measure | Test_method | Group | Comparison | Z | P.unadj | P.adj | Significance |
| --- | --- | --- | --- | --- | --- | --- | --- |
| Acetate | Dunn's Kruskal-Wallis Multiple Comparisons | T0 | T0 - T3 | 1.9984 | 0.0457 | 0.0913 | ns |
| Acetate | Dunn's Kruskal-Wallis Multiple Comparisons | T0 | T0 - T2 | 2.2895 | 0.0221 | 0.0662 | ns |
| Acetate | Dunn's Kruskal-Wallis Multiple Comparisons | T1 | T1 - T0 | -2.3687 | 0.0179 | 0.0714 | ns |
| Acetate | Dunn's Kruskal-Wallis Multiple Comparisons | T1 | T1 - T3 | 3.5682 | 0.0004 | 0.0018 | ** |
| Acetate | Dunn's Kruskal-Wallis Multiple Comparisons | T1 | T1 - T2 | 4.0128 | 0.0001 | 0.0004 | *** |
| Acetate | Dunn's Kruskal-Wallis Multiple Comparisons | T3 | T3 - T2 | 0.1608 | 0.8723 | 0.8723 | ns |
| Ammonium | Dunn's Kruskal-Wallis Multiple Comparisons | T0 | T0 - T3 | 0.2442 | 0.8071 | 0.8071 | ns |
| Ammonium | Dunn's Kruskal-Wallis Multiple Comparisons | T1 | T1 - T2 | 0.5748 | 0.5655 | 1.0000 | ns |
| Ammonium | Dunn's Kruskal-Wallis Multiple Comparisons | T1 | T1 - T0 | -1.3106 | 0.1900 | 1.0000 | ns |
| Ammonium | Dunn's Kruskal-Wallis Multiple Comparisons | T1 | T1 - T3 | 1.2266 | 0.2200 | 1.0000 | ns |
| Ammonium | Dunn's Kruskal-Wallis Multiple Comparisons | T2 | T2 - T0 | -0.7592 | 0.4477 | 1.0000 | ns |
| Ammonium | Dunn's Kruskal-Wallis Multiple Comparisons | T2 | T2 - T3 | 0.7861 | 0.4318 | 1.0000 | ns |
| Calcium | Dunn's Kruskal-Wallis Multiple Comparisons | T0 | T0 - T2 | 0.1493 | 0.8813 | 0.8813 | ns |
| Calcium | Dunn's Kruskal-Wallis Multiple Comparisons | T1 | T1 - T3 | 0.7417 | 0.4583 | 1.0000 | ns |
| Calcium | Dunn's Kruskal-Wallis Multiple Comparisons | T1 | T1 - T0 | -1.4112 | 0.1582 | 0.9491 | ns |
| Calcium | Dunn's Kruskal-Wallis Multiple Comparisons | T1 | T1 - T2 | 1.4042 | 0.1603 | 0.8013 | ns |
| Calcium | Dunn's Kruskal-Wallis Multiple Comparisons | T3 | T3 - T0 | -0.4043 | 0.6860 | 1.0000 | ns |
| Calcium | Dunn's Kruskal-Wallis Multiple Comparisons | T3 | T3 - T2 | -0.4844 | 0.6281 | 1.0000 | ns |
| Chloride | Dunn's Kruskal-Wallis Multiple Comparisons | T0 | T0 - T1 | 0.1628 | 0.8707 | 1.0000 | ns |
| Chloride | Dunn's Kruskal-Wallis Multiple Comparisons | T0 | T0 - T3 | 2.6802 | 0.0074 | 0.0368 | * |
| Chloride | Dunn's Kruskal-Wallis Multiple Comparisons | T0 | T0 - T2 | 3.4691 | 0.0005 | 0.0031 | ** |
| Chloride | Dunn's Kruskal-Wallis Multiple Comparisons | T1 | T1 - T3 | 2.2000 | 0.0278 | 0.0834 | ns |
| Chloride | Dunn's Kruskal-Wallis Multiple Comparisons | T1 | T1 - T2 | 2.6697 | 0.0076 | 0.0304 | * |
| Chloride | Dunn's Kruskal-Wallis Multiple Comparisons | T3 | T3 - T2 | -0.0785 | 0.9375 | 0.9375 | ns |
| CN | Dunn's Kruskal-Wallis Multiple Comparisons | T0 | T0 - T3 | 0.0844 | 0.9327 | 0.9327 | ns |
| CN | Dunn's Kruskal-Wallis Multiple Comparisons | T0 | T0 - T1 | 1.3812 | 0.1672 | 0.8361 | ns |
| CN | Dunn's Kruskal-Wallis Multiple Comparisons | T2 | T2 - T0 | -1.1763 | 0.2395 | 0.9579 | ns |
| CN | Dunn's Kruskal-Wallis Multiple Comparisons | T2 | T2 - T3 | 0.9460 | 0.3441 | 0.6883 | ns |
| CN | Dunn's Kruskal-Wallis Multiple Comparisons | T2 | T2 - T1 | -2.2111 | 0.0270 | 0.1622 | ns |
| CN | Dunn's Kruskal-Wallis Multiple Comparisons | T3 | T3 - T1 | -0.9960 | 0.3192 | 0.9577 | ns |
| d13C | Dunn's Kruskal-Wallis Multiple Comparisons | T0 | T0 - T1 | 1.7168 | 0.0860 | 0.4301 | ns |
| d13C | Dunn's Kruskal-Wallis Multiple Comparisons | T0 | T0 - T2 | 1.1460 | 0.2518 | 0.7554 | ns |
| d13C | Dunn's Kruskal-Wallis Multiple Comparisons | T0 | T0 - T3 | 2.4298 | 0.0151 | 0.0906 | ns |
| d13C | Dunn's Kruskal-Wallis Multiple Comparisons | T1 | T1 - T2 | -0.6298 | 0.5288 | 0.5288 | ns |
| d13C | Dunn's Kruskal-Wallis Multiple Comparisons | T1 | T1 - T3 | 0.7797 | 0.4356 | 0.8712 | ns |
| d13C | Dunn's Kruskal-Wallis Multiple Comparisons | T2 | T2 - T3 | 1.4036 | 0.1604 | 0.6417 | ns |
| d15N | Dunn's Kruskal-Wallis Multiple Comparisons | T0 | T0 - T2 | 1.0153 | 0.3100 | 0.9299 | ns |
| d15N | Dunn's Kruskal-Wallis Multiple Comparisons | T0 | T0 - T1 | 2.3268 | 0.0200 | 0.1198 | ns |
| d15N | Dunn's Kruskal-Wallis Multiple Comparisons | T0 | T0 - T3 | 1.8392 | 0.0659 | 0.3294 | ns |
| d15N | Dunn's Kruskal-Wallis Multiple Comparisons | T1 | T1 - T3 | -0.2052 | 0.8374 | 0.8374 | ns |
| d15N | Dunn's Kruskal-Wallis Multiple Comparisons | T2 | T2 - T1 | -1.2906 | 0.1969 | 0.7874 | ns |
| d15N | Dunn's Kruskal-Wallis Multiple Comparisons | T2 | T2 - T3 | 0.9534 | 0.3404 | 0.6808 | ns |
| Fluoride | Dunn's Kruskal-Wallis Multiple Comparisons | T0 | T0 - T3 | 0.2456 | 0.8060 | 1.0000 | ns |
| Fluoride | Dunn's Kruskal-Wallis Multiple Comparisons | T0 | T0 - T1 | -0.1824 | 0.8553 | 0.8553 | ns |
| Fluoride | Dunn's Kruskal-Wallis Multiple Comparisons | T0 | T0 - T2 | 2.4243 | 0.0153 | 0.0920 | ns |
| Fluoride | Dunn's Kruskal-Wallis Multiple Comparisons | T1 | T1 - T2 | 2.1348 | 0.0328 | 0.1639 | ns |
| Fluoride | Dunn's Kruskal-Wallis Multiple Comparisons | T3 | T3 - T1 | 0.3543 | 0.7231 | 1.0000 | ns |
| Fluoride | Dunn's Kruskal-Wallis Multiple Comparisons | T3 | T3 - T2 | -1.5612 | 0.1185 | 0.4739 | ns |
| Formate | Dunn's Kruskal-Wallis Multiple Comparisons | T0 | T0 - T2 | 2.5438 | 0.0110 | 0.0329 | * |
| Formate | Dunn's Kruskal-Wallis Multiple Comparisons | T0 | T0 - T3 | 3.3312 | 0.0009 | 0.0043 | ** |
| Formate | Dunn's Kruskal-Wallis Multiple Comparisons | T1 | T1 - T0 | -1.1559 | 0.2477 | 0.2477 | ns |
| Formate | Dunn's Kruskal-Wallis Multiple Comparisons | T1 | T1 - T2 | 3.1170 | 0.0018 | 0.0073 | ** |
| Formate | Dunn's Kruskal-Wallis Multiple Comparisons | T1 | T1 - T3 | 3.7859 | 0.0002 | 0.0009 | *** |
| Formate | Dunn's Kruskal-Wallis Multiple Comparisons | T2 | T2 - T3 | 1.2067 | 0.2275 | 0.4551 | ns |
| Fumarate | Dunn's Kruskal-Wallis Multiple Comparisons | T0 | T0 - T1 | 1.3348 | 0.1819 | 0.7277 | ns |
| Fumarate | Dunn's Kruskal-Wallis Multiple Comparisons | T0 | T0 - T2 | 2.5386 | 0.0111 | 0.0668 | ns |
| Fumarate | Dunn's Kruskal-Wallis Multiple Comparisons | T0 | T0 - T3 | 2.3046 | 0.0212 | 0.1059 | ns |
| Fumarate | Dunn's Kruskal-Wallis Multiple Comparisons | T1 | T1 - T2 | 0.8485 | 0.3962 | 0.7923 | ns |
| Fumarate | Dunn's Kruskal-Wallis Multiple Comparisons | T1 | T1 - T3 | 0.9667 | 0.3337 | 1.0000 | ns |
| Fumarate | Dunn's Kruskal-Wallis Multiple Comparisons | T2 | T2 - T3 | 0.2604 | 0.7946 | 0.7946 | ns |
| Lactate | Dunn's Kruskal-Wallis Multiple Comparisons | T0 | T0 - T3 | 0.3977 | 0.6908 | 0.6908 | ns |
| Lactate | Dunn's Kruskal-Wallis Multiple Comparisons | T1 | T1 - T0 | -0.6566 | 0.5115 | 1.0000 | ns |
| Lactate | Dunn's Kruskal-Wallis Multiple Comparisons | T1 | T1 - T3 | 0.8535 | 0.3934 | 1.0000 | ns |
| Lactate | Dunn's Kruskal-Wallis Multiple Comparisons | T2 | T2 - T1 | -0.7874 | 0.4310 | 1.0000 | ns |
| Lactate | Dunn's Kruskal-Wallis Multiple Comparisons | T2 | T2 - T0 | -1.7043 | 0.0883 | 0.5300 | ns |
| Lactate | Dunn's Kruskal-Wallis Multiple Comparisons | T2 | T2 - T3 | 1.6255 | 0.1041 | 0.5203 | ns |
| Magnesium | Dunn's Kruskal-Wallis Multiple Comparisons | T0 | T0 - T2 | 1.9354 | 0.0529 | 0.2647 | ns |
| Magnesium | Dunn's Kruskal-Wallis Multiple Comparisons | T0 | T0 - T3 | 2.0464 | 0.0407 | 0.2443 | ns |
| Magnesium | Dunn's Kruskal-Wallis Multiple Comparisons | T1 | T1 - T0 | -0.1388 | 0.8896 | 0.8896 | ns |
| Magnesium | Dunn's Kruskal-Wallis Multiple Comparisons | T1 | T1 - T2 | 1.6982 | 0.0895 | 0.2684 | ns |
| Magnesium | Dunn's Kruskal-Wallis Multiple Comparisons | T1 | T1 - T3 | 1.8834 | 0.0596 | 0.2386 | ns |
| Magnesium | Dunn's Kruskal-Wallis Multiple Comparisons | T2 | T2 - T3 | 0.4663 | 0.6410 | 1.0000 | ns |
| Nitrate | Dunn's Kruskal-Wallis Multiple Comparisons | T0 | T0 - T3 | 1.3025 | 0.1927 | 0.3855 | ns |
| Nitrate | Dunn's Kruskal-Wallis Multiple Comparisons | T1 | T1 - T0 | -1.5645 | 0.1177 | 0.3531 | ns |
| Nitrate | Dunn's Kruskal-Wallis Multiple Comparisons | T1 | T1 - T3 | 2.3416 | 0.0192 | 0.0960 | ns |
| Nitrate | Dunn's Kruskal-Wallis Multiple Comparisons | T2 | T2 - T1 | -0.4005 | 0.6888 | 0.6888 | ns |
| Nitrate | Dunn's Kruskal-Wallis Multiple Comparisons | T2 | T2 - T0 | -2.2441 | 0.0248 | 0.0993 | ns |
| Nitrate | Dunn's Kruskal-Wallis Multiple Comparisons | T2 | T2 - T3 | 2.8612 | 0.0042 | 0.0253 | * |
| Nitrite | Dunn's Kruskal-Wallis Multiple Comparisons | T0 | T0 - T1 | -1.1204 | 0.2625 | 1.0000 | ns |
| Nitrite | Dunn's Kruskal-Wallis Multiple Comparisons | T2 | T2 - T0 | -1.7494 | 0.0802 | 0.4011 | ns |
| Nitrite | Dunn's Kruskal-Wallis Multiple Comparisons | T2 | T2 - T1 | -0.4024 | 0.6874 | 1.0000 | ns |
| Nitrite | Dunn's Kruskal-Wallis Multiple Comparisons | T3 | T3 - T2 | -0.3700 | 0.7114 | 0.7114 | ns |
| Nitrite | Dunn's Kruskal-Wallis Multiple Comparisons | T3 | T3 - T0 | -1.7940 | 0.0728 | 0.4369 | ns |
| Nitrite | Dunn's Kruskal-Wallis Multiple Comparisons | T3 | T3 - T1 | -0.6896 | 0.4905 | 1.0000 | ns |
| Oxalate | Dunn's Kruskal-Wallis Multiple Comparisons | T0 | T0 - T2 | 1.5879 | 0.1123 | 0.3370 | ns |
| Oxalate | Dunn's Kruskal-Wallis Multiple Comparisons | T0 | T0 - T3 | 2.6533 | 0.0080 | 0.0399 | * |
| Oxalate | Dunn's Kruskal-Wallis Multiple Comparisons | T1 | T1 - T0 | -0.8811 | 0.3783 | 0.3783 | ns |
| Oxalate | Dunn's Kruskal-Wallis Multiple Comparisons | T1 | T1 - T2 | 2.0907 | 0.0366 | 0.1462 | ns |
| Oxalate | Dunn's Kruskal-Wallis Multiple Comparisons | T1 | T1 - T3 | 2.9848 | 0.0028 | 0.0170 | * |
| Oxalate | Dunn's Kruskal-Wallis Multiple Comparisons | T2 | T2 - T3 | 1.2845 | 0.1990 | 0.3980 | ns |
| pH | Dunn's Kruskal-Wallis Multiple Comparisons | T1 | T1 - T2 | 0.9174 | 0.3590 | 0.7179 | ns |
| pH | Dunn's Kruskal-Wallis Multiple Comparisons | T1 | T1 - T3 | 1.5580 | 0.1192 | 0.4769 | ns |
| pH | Dunn's Kruskal-Wallis Multiple Comparisons | T1 | T1 - T0 | -3.5920 | 0.0003 | 0.0020 | ** |
| pH | Dunn's Kruskal-Wallis Multiple Comparisons | T2 | T2 - T3 | 0.8285 | 0.4074 | 0.4074 | ns |
| pH | Dunn's Kruskal-Wallis Multiple Comparisons | T2 | T2 - T0 | -2.8908 | 0.0038 | 0.0192 | * |
| pH | Dunn's Kruskal-Wallis Multiple Comparisons | T3 | T3 - T0 | -1.4090 | 0.1588 | 0.4765 | ns |
| Phosphate | Dunn's Kruskal-Wallis Multiple Comparisons | T0 | T0 - T3 | 1.3846 | 0.1662 | 0.9971 | ns |
| Phosphate | Dunn's Kruskal-Wallis Multiple Comparisons | T0 | T0 - T1 | 1.2709 | 0.2038 | 1.0000 | ns |
| Phosphate | Dunn's Kruskal-Wallis Multiple Comparisons | T0 | T0 - T2 | 1.1751 | 0.2400 | 0.9598 | ns |
| Phosphate | Dunn's Kruskal-Wallis Multiple Comparisons | T1 | T1 - T2 | -0.2009 | 0.8408 | 0.8408 | ns |
| Phosphate | Dunn's Kruskal-Wallis Multiple Comparisons | T3 | T3 - T1 | 0.2177 | 0.8276 | 1.0000 | ns |
| Phosphate | Dunn's Kruskal-Wallis Multiple Comparisons | T3 | T3 - T2 | 0.4147 | 0.6784 | 1.0000 | ns |
| Potassium | Dunn's Kruskal-Wallis Multiple Comparisons | T1 | T1 - T3 | -0.6995 | 0.4842 | 0.4842 | ns |
| Potassium | Dunn's Kruskal-Wallis Multiple Comparisons | T1 | T1 - T2 | 1.0036 | 0.3156 | 0.6311 | ns |
| Potassium | Dunn's Kruskal-Wallis Multiple Comparisons | T1 | T1 - T0 | -2.3628 | 0.0181 | 0.0907 | ns |
| Potassium | Dunn's Kruskal-Wallis Multiple Comparisons | T2 | T2 - T0 | -1.4089 | 0.1589 | 0.4766 | ns |
| Potassium | Dunn's Kruskal-Wallis Multiple Comparisons | T3 | T3 - T2 | -1.6576 | 0.0974 | 0.3896 | ns |
| Potassium | Dunn's Kruskal-Wallis Multiple Comparisons | T3 | T3 - T0 | -2.9138 | 0.0036 | 0.0214 | * |
| Pyruvate | Dunn's Kruskal-Wallis Multiple Comparisons | T0 | T0 - T3 | 0.3965 | 0.6917 | 1.0000 | ns |
| Pyruvate | Dunn's Kruskal-Wallis Multiple Comparisons | T1 | T1 - T0 | -0.2796 | 0.7798 | 0.7798 | ns |
| Pyruvate | Dunn's Kruskal-Wallis Multiple Comparisons | T1 | T1 - T3 | 0.5605 | 0.5751 | 1.0000 | ns |
| Pyruvate | Dunn's Kruskal-Wallis Multiple Comparisons | T2 | T2 - T1 | -1.3644 | 0.1724 | 0.6897 | ns |
| Pyruvate | Dunn's Kruskal-Wallis Multiple Comparisons | T2 | T2 - T0 | -1.9927 | 0.0463 | 0.2777 | ns |
| Pyruvate | Dunn's Kruskal-Wallis Multiple Comparisons | T2 | T2 - T3 | 1.8372 | 0.0662 | 0.3309 | ns |
| Sodium | Dunn's Kruskal-Wallis Multiple Comparisons | T0 | T0 - T3 | 0.5100 | 0.6101 | 0.6101 | ns |
| Sodium | Dunn's Kruskal-Wallis Multiple Comparisons | T0 | T0 - T2 | 3.1413 | 0.0017 | 0.0084 | ** |
| Sodium | Dunn's Kruskal-Wallis Multiple Comparisons | T1 | T1 - T0 | -1.1512 | 0.2496 | 0.4993 | ns |
| Sodium | Dunn's Kruskal-Wallis Multiple Comparisons | T1 | T1 - T3 | 1.3339 | 0.1822 | 0.5467 | ns |
| Sodium | Dunn's Kruskal-Wallis Multiple Comparisons | T1 | T1 - T2 | 3.5980 | 0.0003 | 0.0019 | ** |
| Sodium | Dunn's Kruskal-Wallis Multiple Comparisons | T3 | T3 - T2 | -1.8454 | 0.0650 | 0.2599 | ns |
| Succinate | Dunn's Kruskal-Wallis Multiple Comparisons | T0 | T0 - T2 | -0.2781 | 0.7809 | 0.7809 | ns |
| Succinate | Dunn's Kruskal-Wallis Multiple Comparisons | T0 | T0 - T3 | 0.7206 | 0.4711 | 1.0000 | ns |
| Succinate | Dunn's Kruskal-Wallis Multiple Comparisons | T1 | T1 - T0 | -0.6395 | 0.5225 | 1.0000 | ns |
| Succinate | Dunn's Kruskal-Wallis Multiple Comparisons | T1 | T1 - T2 | 0.3554 | 0.7223 | 1.0000 | ns |
| Succinate | Dunn's Kruskal-Wallis Multiple Comparisons | T1 | T1 - T3 | 1.1205 | 0.2625 | 1.0000 | ns |
| Succinate | Dunn's Kruskal-Wallis Multiple Comparisons | T2 | T2 - T3 | 0.8722 | 0.3831 | 1.0000 | ns |
| Sulfate | Dunn's Kruskal-Wallis Multiple Comparisons | T0 | T0 - T3 | 1.6061 | 0.1083 | 0.5413 | ns |
| Sulfate | Dunn's Kruskal-Wallis Multiple Comparisons | T1 | T1 - T2 | 1.4238 | 0.1545 | 0.3090 | ns |
| Sulfate | Dunn's Kruskal-Wallis Multiple Comparisons | T1 | T1 - T0 | -1.5027 | 0.1329 | 0.5317 | ns |
| Sulfate | Dunn's Kruskal-Wallis Multiple Comparisons | T1 | T1 - T3 | 2.5572 | 0.0106 | 0.0633 | ns |
| Sulfate | Dunn's Kruskal-Wallis Multiple Comparisons | T2 | T2 - T0 | 0.0711 | 0.9433 | 0.9433 | ns |
| Sulfate | Dunn's Kruskal-Wallis Multiple Comparisons | T2 | T2 - T3 | 1.4342 | 0.1515 | 0.4546 | ns |
| TC | Dunn's Kruskal-Wallis Multiple Comparisons | T0 | T0 - T1 | 0.7489 | 0.4539 | 0.9079 | ns |
| TC | Dunn's Kruskal-Wallis Multiple Comparisons | T2 | T2 - T3 | 0.1657 | 0.8684 | 0.8684 | ns |
| TC | Dunn's Kruskal-Wallis Multiple Comparisons | T2 | T2 - T0 | -1.5833 | 0.1134 | 0.5668 | ns |
| TC | Dunn's Kruskal-Wallis Multiple Comparisons | T2 | T2 - T1 | -1.9668 | 0.0492 | 0.2953 | ns |
| TC | Dunn's Kruskal-Wallis Multiple Comparisons | T3 | T3 - T0 | -1.0829 | 0.2788 | 0.8365 | ns |
| TC | Dunn's Kruskal-Wallis Multiple Comparisons | T3 | T3 - T1 | -1.5196 | 0.1286 | 0.5144 | ns |
| TN | Dunn's Kruskal-Wallis Multiple Comparisons | T0 | T0 - T1 | 0.7126 | 0.4761 | 0.9521 | ns |
| TN | Dunn's Kruskal-Wallis Multiple Comparisons | T2 | T2 - T0 | -0.7645 | 0.4446 | 1.0000 | ns |
| TN | Dunn's Kruskal-Wallis Multiple Comparisons | T2 | T2 - T1 | -1.2688 | 0.2045 | 1.0000 | ns |
| TN | Dunn's Kruskal-Wallis Multiple Comparisons | T3 | T3 - T2 | -0.3448 | 0.7303 | 0.7303 | ns |
| TN | Dunn's Kruskal-Wallis Multiple Comparisons | T3 | T3 - T0 | -0.9818 | 0.3262 | 1.0000 | ns |
| TN | Dunn's Kruskal-Wallis Multiple Comparisons | T3 | T3 - T1 | -1.4038 | 0.1604 | 0.9622 | ns |

Table T 2 shows the Letter Display output for the multiple comparison significance testing on environmental variables. Different letters represent significant differences.

| **Measure** | **Test_method** | **Group** | **Letter** | **MonoLetter** |
| --- | --- | --- | --- | --- |
| Acetate | Dunn's Kruskal-Wallis Multiple Comparisons | T0 | ab | ab |
| Acetate | Dunn's Kruskal-Wallis Multiple Comparisons | T1 | a | a |
| Acetate | Dunn's Kruskal-Wallis Multiple Comparisons | T2 | b | b |
| Acetate | Dunn's Kruskal-Wallis Multiple Comparisons | T3 | b | b |
| Ammonium | Dunn's Kruskal-Wallis Multiple Comparisons | T0 | a | a |
| Ammonium | Dunn's Kruskal-Wallis Multiple Comparisons | T1 | a | a |
| Ammonium | Dunn's Kruskal-Wallis Multiple Comparisons | T2 | a | a |
| Ammonium | Dunn's Kruskal-Wallis Multiple Comparisons | T3 | a | a |
| Calcium | Dunn's Kruskal-Wallis Multiple Comparisons | T0 | a | a |
| Calcium | Dunn's Kruskal-Wallis Multiple Comparisons | T1 | a | a |
| Calcium | Dunn's Kruskal-Wallis Multiple Comparisons | T2 | a | a |
| Calcium | Dunn's Kruskal-Wallis Multiple Comparisons | T3 | a | a |
| Chloride | Dunn's Kruskal-Wallis Multiple Comparisons | T0 | a | a |
| Chloride | Dunn's Kruskal-Wallis Multiple Comparisons | T1 | ab | ab |
| Chloride | Dunn's Kruskal-Wallis Multiple Comparisons | T2 | c | c |
| Chloride | Dunn's Kruskal-Wallis Multiple Comparisons | T3 | bc | bc |
| CN | Dunn's Kruskal-Wallis Multiple Comparisons | T0 | a | a |
| CN | Dunn's Kruskal-Wallis Multiple Comparisons | T1 | a | a |
| CN | Dunn's Kruskal-Wallis Multiple Comparisons | T2 | a | a |
| CN | Dunn's Kruskal-Wallis Multiple Comparisons | T3 | a | a |
| d13C | Dunn's Kruskal-Wallis Multiple Comparisons | T0 | a | a |
| d13C | Dunn's Kruskal-Wallis Multiple Comparisons | T1 | a | a |
| d13C | Dunn's Kruskal-Wallis Multiple Comparisons | T2 | a | a |
| d13C | Dunn's Kruskal-Wallis Multiple Comparisons | T3 | a | a |
| d15N | Dunn's Kruskal-Wallis Multiple Comparisons | T0 | a | a |
| d15N | Dunn's Kruskal-Wallis Multiple Comparisons | T1 | a | a |
| d15N | Dunn's Kruskal-Wallis Multiple Comparisons | T2 | a | a |
| d15N | Dunn's Kruskal-Wallis Multiple Comparisons | T3 | a | a |
| Fluoride | Dunn's Kruskal-Wallis Multiple Comparisons | T0 | a | a |
| Fluoride | Dunn's Kruskal-Wallis Multiple Comparisons | T1 | a | a |
| Fluoride | Dunn's Kruskal-Wallis Multiple Comparisons | T2 | a | a |
| Fluoride | Dunn's Kruskal-Wallis Multiple Comparisons | T3 | a | a |
| Formate | Dunn's Kruskal-Wallis Multiple Comparisons | T0 | a | a |
| Formate | Dunn's Kruskal-Wallis Multiple Comparisons | T1 | a | a |
| Formate | Dunn's Kruskal-Wallis Multiple Comparisons | T2 | b | b |
| Formate | Dunn's Kruskal-Wallis Multiple Comparisons | T3 | b | b |
| Fumarate | Dunn's Kruskal-Wallis Multiple Comparisons | T0 | a | a |
| Fumarate | Dunn's Kruskal-Wallis Multiple Comparisons | T1 | a | a |
| Fumarate | Dunn's Kruskal-Wallis Multiple Comparisons | T2 | a | a |
| Fumarate | Dunn's Kruskal-Wallis Multiple Comparisons | T3 | a | a |
| Lactate | Dunn's Kruskal-Wallis Multiple Comparisons | T0 | a | a |
| Lactate | Dunn's Kruskal-Wallis Multiple Comparisons | T1 | a | a |
| Lactate | Dunn's Kruskal-Wallis Multiple Comparisons | T2 | a | a |
| Lactate | Dunn's Kruskal-Wallis Multiple Comparisons | T3 | a | a |
| Magnesium | Dunn's Kruskal-Wallis Multiple Comparisons | T0 | a | a |
| Magnesium | Dunn's Kruskal-Wallis Multiple Comparisons | T1 | a | a |
| Magnesium | Dunn's Kruskal-Wallis Multiple Comparisons | T2 | a | a |
| Magnesium | Dunn's Kruskal-Wallis Multiple Comparisons | T3 | a | a |
| Nitrate | Dunn's Kruskal-Wallis Multiple Comparisons | T0 | ab | ab |
| Nitrate | Dunn's Kruskal-Wallis Multiple Comparisons | T1 | ab | ab |
| Nitrate | Dunn's Kruskal-Wallis Multiple Comparisons | T2 | a | a |
| Nitrate | Dunn's Kruskal-Wallis Multiple Comparisons | T3 | b | b |
| Nitrite | Dunn's Kruskal-Wallis Multiple Comparisons | T0 | a | a |
| Nitrite | Dunn's Kruskal-Wallis Multiple Comparisons | T1 | a | a |
| Nitrite | Dunn's Kruskal-Wallis Multiple Comparisons | T2 | a | a |
| Nitrite | Dunn's Kruskal-Wallis Multiple Comparisons | T3 | a | a |
| Oxalate | Dunn's Kruskal-Wallis Multiple Comparisons | T0 | a | a |
| Oxalate | Dunn's Kruskal-Wallis Multiple Comparisons | T1 | a | a |
| Oxalate | Dunn's Kruskal-Wallis Multiple Comparisons | T2 | ab | ab |
| Oxalate | Dunn's Kruskal-Wallis Multiple Comparisons | T3 | b | b |
| pH | Dunn's Kruskal-Wallis Multiple Comparisons | T0 | b | b |
| pH | Dunn's Kruskal-Wallis Multiple Comparisons | T1 | a | a |
| pH | Dunn's Kruskal-Wallis Multiple Comparisons | T2 | a | a |
| pH | Dunn's Kruskal-Wallis Multiple Comparisons | T3 | ab | ab |
| Phosphate | Dunn's Kruskal-Wallis Multiple Comparisons | T0 | a | a |
| Phosphate | Dunn's Kruskal-Wallis Multiple Comparisons | T1 | a | a |
| Phosphate | Dunn's Kruskal-Wallis Multiple Comparisons | T2 | a | a |
| Phosphate | Dunn's Kruskal-Wallis Multiple Comparisons | T3 | a | a |
| Potassium | Dunn's Kruskal-Wallis Multiple Comparisons | T0 | b | b |
| Potassium | Dunn's Kruskal-Wallis Multiple Comparisons | T1 | ab | ab |
| Potassium | Dunn's Kruskal-Wallis Multiple Comparisons | T2 | ab | ab |
| Potassium | Dunn's Kruskal-Wallis Multiple Comparisons | T3 | a | a |
| Pyruvate | Dunn's Kruskal-Wallis Multiple Comparisons | T0 | a | a |
| Pyruvate | Dunn's Kruskal-Wallis Multiple Comparisons | T1 | a | a |
| Pyruvate | Dunn's Kruskal-Wallis Multiple Comparisons | T2 | a | a |
| Pyruvate | Dunn's Kruskal-Wallis Multiple Comparisons | T3 | a | a |
| Sodium | Dunn's Kruskal-Wallis Multiple Comparisons | T0 | a | a |
| Sodium | Dunn's Kruskal-Wallis Multiple Comparisons | T1 | a | a |
| Sodium | Dunn's Kruskal-Wallis Multiple Comparisons | T2 | b | b |
| Sodium | Dunn's Kruskal-Wallis Multiple Comparisons | T3 | ab | ab |
| Succinate | Dunn's Kruskal-Wallis Multiple Comparisons | T0 | a | a |
| Succinate | Dunn's Kruskal-Wallis Multiple Comparisons | T1 | a | a |
| Succinate | Dunn's Kruskal-Wallis Multiple Comparisons | T2 | a | a |
| Succinate | Dunn's Kruskal-Wallis Multiple Comparisons | T3 | a | a |
| Sulfate | Dunn's Kruskal-Wallis Multiple Comparisons | T0 | a | a |
| Sulfate | Dunn's Kruskal-Wallis Multiple Comparisons | T1 | a | a |
| Sulfate | Dunn's Kruskal-Wallis Multiple Comparisons | T2 | a | a |
| Sulfate | Dunn's Kruskal-Wallis Multiple Comparisons | T3 | a | a |
| TC | Dunn's Kruskal-Wallis Multiple Comparisons | T0 | a | a |
| TC | Dunn's Kruskal-Wallis Multiple Comparisons | T1 | a | a |
| TC | Dunn's Kruskal-Wallis Multiple Comparisons | T2 | a | a |
| TC | Dunn's Kruskal-Wallis Multiple Comparisons | T3 | a | a |
| TN | Dunn's Kruskal-Wallis Multiple Comparisons | T0 | a | a |
| TN | Dunn's Kruskal-Wallis Multiple Comparisons | T1 | a | a |
| TN | Dunn's Kruskal-Wallis Multiple Comparisons | T2 | a | a |
| TN | Dunn's Kruskal-Wallis Multiple Comparisons | T3 | a | a |
